# Dynamic Hierarchical Interleaved Bloom Filter: An Updatable Index for Large-Scale Fast Sequence Search

**DOI:** 10.64898/2026.08.26.747224

**Authors:** Enrico Seiler, Myrthe Willemsen, Vitor C. Piro, Knut Reinert

**Author notes:** Equal contributor (Enrico Seiler, Myrthe Willemsen).

## Abstract

**Motivation:** A continued decrease in sequencing costs has facilitated the exponential increase in available sequencing data, with public databases like the European Nucleotide Archive (ENA) and Sequence Read Archive (SRA) reaching well in the order of petabases [1, 2]. This has been the incentive to develop more scalable tools for common bioinformatics tasks. One such task is the approximate searching of short sequence patterns like genes or reads in reference data sets. In recent years, a variety of indexing data structures have been proposed for searching large sequencing databases. The state-of-the-art index, the Hierarchical Interleaved Bloom Filter (HIBF), was first-in-class to index one million samples. However, its static design necessitates costly full reconstructions as repositories grow, motivating an extension to support dynamic updates.

**Results:** In this paper, we introduce a scalable and updatable sequence-search index by extending the HIBF with partial rebuilding to support efficient updates. We demonstrate the Dynamic HIBF’s capacity for large-scale data by iteratively creating an index from over 100 TB of compressed reads across more than 39,000 full human RNA-Seq samples, updated in consecutive batches of 100. To benchmark against state-of-the-art tools, we evaluated incremental performance on a subset of 5,000 samples sub-sampled to 1 % of their original read depth. In this comparative setting, the dynamic HIBF completed the sequential insertion of all 5,000 samples in 5.3 hours—23 to 42 times faster than competing methods and twice as fast as the static HIBF.

## 1 Introduction

The desire to unravel the blueprint of life once inspired the launch of the human genome project in 1990 [3, 4]. The highly ambitious undertaking became an accelerator for the innovation of high-throughput sequencing technologies. The 1000 Genomes Project in 2008 [5], and the subsequent 100,000 Genomes Project in 2014 [6], have steered the continued development in next- and later-generation sequencing. Within two decades, the throughput of a single machine increased dramatically from 21 billion base pairs collected over months to multiple terabases in a couple of hours [7, 8]. As a consequence, sequencing costs for entire human genomes have dropped well to as low as 100 USD per sample, making the generation of data more accessible. DNA, RNA, and even protein sequencing experiments have become an integral part of a multitude of biological research areas.

This Big Data era has fueled the exponential rise in available sequencing data, among private repositories and public databases. For instance, the raw sequencing experiment data sets like the Sequence Read Archive (SRA) comprise more than 28 Pebibytes (PiB) of nucleotide sequences [1]. The European Nucleotide Archive (ENA) has grown to around 99 PiB of sequences [2]. Studies by international consortia, including The Cancer Genome Atlas (TCGA), the International Cancer Genomics Consortium (ICGC), the Genotype-Tissue Expression (GTEx) project, and the Metagenomics & Metadesign of Subways & Urban Biomes (MetaSUB) undertaking, generated hundreds of terabytes of data and continue to grow with new samples added on a daily basis.

These large sequence repositories collectively possess the statistical power to answer biomedical research questions previously unanswerable and trigger the development of novel diagnoses and treatments. They further improve our understanding of ecology and microbiology and contribute to sustainable biotechnological innovations. However, accessing the entirety of the data at once is close to impossible, limiting scientific discovery opportunities. Even for defined subsets, it can be complicated to access the data, i.e., transferring, storing, and searching it, let alone any of the more complex downstream analyses, including haplotyping, *de novo* assembly, variant calling, error correction, read mapping, and transcript quantification, to name a few. Accessing larger amounts of data demands extensive computational resources, restraining the data to privileged research labs only.

The need for scalability has been the incentive to develop more efficient tools for various bioinformatics tasks. One such task, referred to as multiset-membership query, involves searching for short sequence patterns in reference data sets. The objective is to identify which datasets contain the exact or a similar pattern. This task has broad applications and serves as the basis for numerous downstream investigations. The naive approach to querying sequentially compares the query sequence against every position in each reference dataset. However, this approach is not scalable to large datasets due to high computational cost.

To enable more efficient queries, additional data structures, referred to as *indexes*, have been developed on top of the sequencing databases. Many indexes have been designed and implemented in a dozen of tools over the last decades, summarized in Section 1.3.

### 1.1 Biological applications

The task of multiset-membership queries has found diverse applications in disciplines dealing with processing and analysing large bodies of text.

A common application of indexes is to identify species by the DNA collected from environmental samples or clinical infection studies. Depending on the research focus, a specific taxonomic domain could be selected as reference datasets, for instance, RefSeq Bacteria [9] for microbial communities or RefSeq Virus for studying viral infections. Alternatively, broader databases, like the Sequence Read Archive (SRA), can be used for exploring diverse samples and open research questions. SRA grows by more than a thousand terabases per year, and regular updates will be necessary to incorporate those new experiments. When sets represent single accessions or experiments, the majority of these updates will be insertions of new sets (samples). If instead, sets represent species, also set modifications or deletions may be desired to remove false data points [10].

Secondly, an index can be used as a prefilter for more advanced analyses, such as exact read mapping. By querying reads on the index first, it becomes possible to efficiently distribute the computational load of working with large full-text indices like FM-indices or suffix arrays.

A third application is genotyping, which refers to measuring variants of single nucleotide polymorphisms (SNPs) and small insertions and deletions (indels) in sequencing data. The collections of SNPs and indels for individual humans can be indexed as separate sets. Alternatively, the SNPs in the NCBI’s dbSNP [11] database can be categorized by associated phenotype and indexed as such. The approximate membership queries can be leveraged to efficiently detect if SNPs in new samples are linked to certain disease conditions.

Further applications of indexes consist of profiling of functional alterations in RNA-seq data, exploration of gene fusions, and quantification of RNA transcripts per gene.

### 1.2 Dynamization methods

*Dynamization* refers to the process of making a data structure dynamic. Generally, data structures are categorized as dynamic when they allow changing the underlying input set through inserting or deleting items. In the context of indexes that store multiple sequence sets, dynamic operations include inserting, deleting, and modifying sets.

While many approaches for dynamic updates exist (see Section 1.3), existing implementations exhibit six primary limitations in practice:

1. Prohibitive preprocessing requirements (e.g., computationally expensive *k*-mer counting)
2. Codebases functioning primarily as proofs-of-concept rather than production-ready software
3. Software instability and execution failures on large-scale datasets
4. Lack of active software maintenance and long-term support
5. Suboptimal empirical scalability across index construction, updating, **and** querying
6. Computationally inefficient update mechanisms, often necessitating partial or full index reconstruction

Dynamization strategies can be broadly categorized into two classes. The first class involves managing updates using a “smaller” auxiliary data structure compared to the main data structure. One example of such a strategy is the Bentley-Saxe transformation [12], which can be applied to static data structures that support efficient merges. In this approach, the data elements are maintained in a set of static data structures with varying sizes. The number of these structures scales with the logarithm of the number of elements.

The second class of dynamization strategies does not rely on additional data structures. Instead, insertions and deletions are handled by local or global modifications directly within the structure itself, for instance, by reorganizing the data sets or the local building blocks of the index. In 1983, Overmars described various techniques for reorganizing data structures, including local, partial, and global rebuilding [13].

*Local rebuilding*, also known as balancing, involves making minor adjustments to ensure that the data remains properly distributed after each update. *Partial rebuilding* is a technique that periodically rebuilds only the degenerated parts of the data structure, restoring a balanced state.

*Global rebuilding* refers to the complete reconstruction of the entire structure. While this approach may be suitable for applications with frequent simultaneous updates, it is costly. Data structures that rely solely on global rebuilding for updates are typically not regarded as dynamic unless the rebuilding cost can be amortized. While these techniques are applicable in a generic sense, it is important to note that most dynamization methods are tailored to specific data structures or search problems that satisfy certain properties or constraints. It may not be feasible to devise a universal methodology that achieves optimal performance across all costs [14]. Designing efficient dynamization methods for complex data structures often necessitates careful engineering and thoughtful evaluation of the theoretical and actual resource usage. As a result, many indexes rely on expensive global rebuilding without incorporating suitable update mechanisms.

### 1.3 Multiset indexes

Many data structures for storing multiple sequencing sets, or multisets, have been developed to provide an answer to the experiment discovery problem. An overview of various indexes is shown in Table 1 and is further elaborated on below. Table 1 also shows which dynamic operations the indexes can perform. An expanded version of this table, featuring more tools, can be found in Appendix Table 3. Some indexes also have additional functionalities, such as counting, positional information, or alignments. A more detailed discussion on indexing multisets can be found in the review by Marchet *et al*. [15].

**Table 1.**
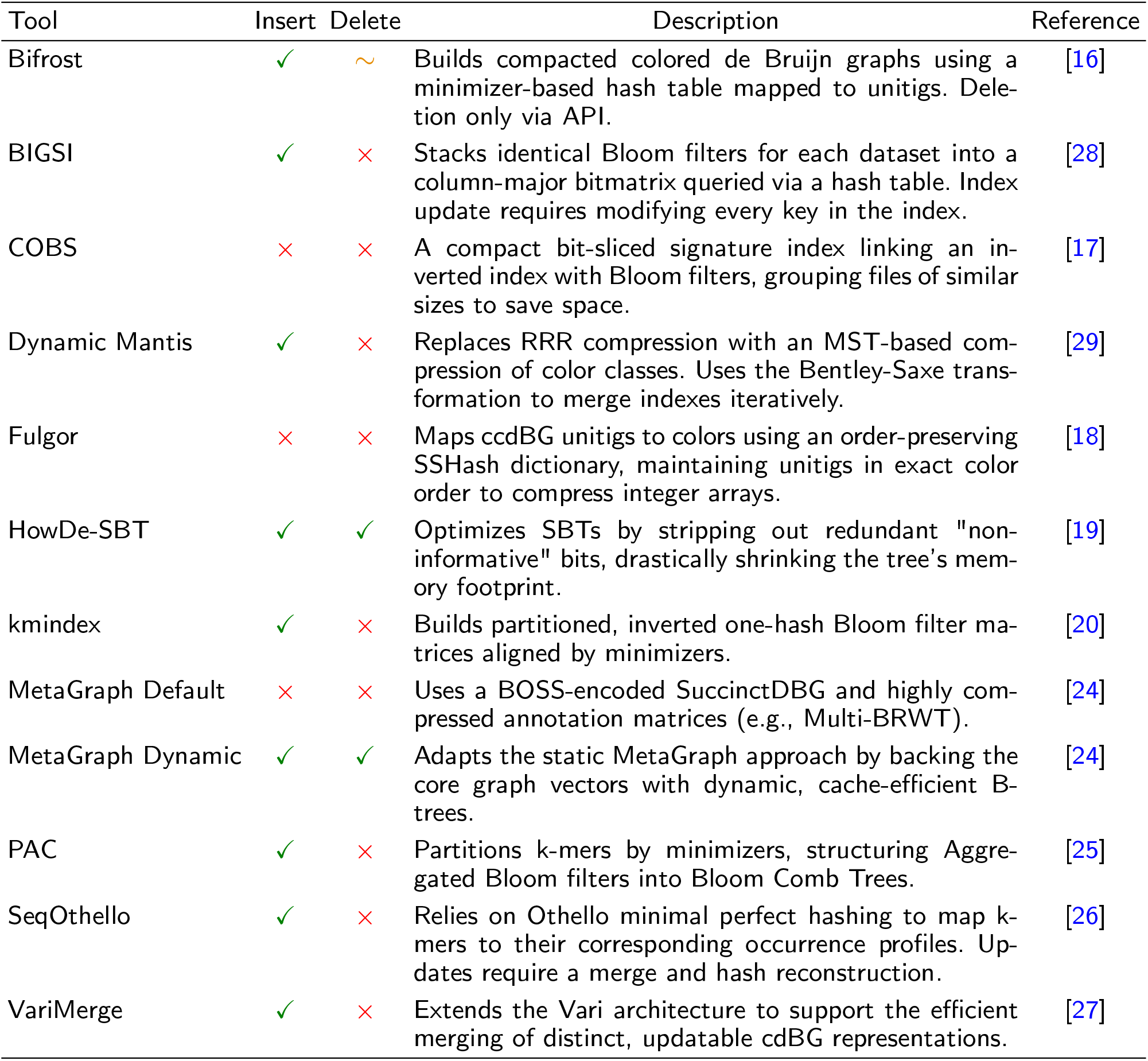
Algorithmic comparison of multi-set sequence indexing tools. Overview of representative colored de Bruijn graph and sequence aggregation indices, detailing their support for dynamic insertion and deletion operations along with their core indexing mechanisms. An expanded version of this table can be found in Appendix Table 3.

**Bifrost** [16] is a representation of a colored de Bruijn graph with minimizers and blocked Bloom filters. A hash function is used to associate *k*-mers with the right BF. Bifrost supports balanced insertions of new sequences, although it does not describe how it deals with the increasing false positive rate.

**COBS** [17], the COmpact Bit-sliced Signature index, improves the framework of BIGSI by allowing for variable-length BFs to adapt for varying data set sizes. It stores similar-sized *k*-mer sets in blocks of equally-sized bloom filters within a matrix. To provide more scalability, COBS is designed such that it does not need the complete index in RAM, but does not support updates.

**Fulgor** [18] maps ccdBG unitigs to colors using an order-preserving SSHash dictionary, maintaining unitigs in exact color order to heavily compress integer arrays. The strict unitig ordering and immutable succinct structures mandate total reconstruction for updates.

The **Hierarchical Interleaved Bloom Filters** (HIBF), which we address in this paper, is a hierarchical structure of multiple Interleaved Bloom Filter (IBF). The HIBF uncouples the *k*-mer sets provided by the user from the separate bloom filters stored within the IBF to enable the indexing of larger datasets with varying sample sizes. A detailed description can be found in Section 2.1.

**HowDeSBT** [19] optimizes the Sequence Bloom Tree (SBT) by stripping out redundant, non-informative bits, drastically shrinking the tree’s memory footprint. The tree topology natively enables dynamic updates.

**kmindex** [20] builds partitioned, inverted one-hash Bloom filter matrices aligned by minimizers, leaving empty padding bits per row. Builds upon the kmtricks pipeline [21]. Datasets can be seamlessly appended, but sequence deletion is currently impossible.

**Mantis** [22] is based on Counting Quotient Filters and associates each *k*-mer with a color ID, an identifier of a specific combination of data sets. Each color ID, in turn, is used to obtain color-class bit vectors from the color-class matrix, indicating in which sets a queried *k*-mer occurs.

**Dynamic Mantis** [23] is an extension of Mantis. It uses minimum spanning trees (MST) to encode similar sequence sets sharing *k*-mers more efficiently. The data structure was further extended to support insertions of new sets by using the Bentley-Saxe transformation.

**MetaGraph** [24] is an index employing annotated node-centric de Bruijn graphs. For each set, a separate de Bruijn graph is constructed. These dBGs are then cleaned, merged, and stored as a succinct BOSS table. An annotation matrix is used to associate the *k*-mers in the dBG with the labels of its originating sequences. The MetaGraph framework provides a wide range of data structures for various purposes, including dynamic updates.

**PAC** [25] partitions k-mers by minimizers, and structures Aggregated Bloom filters into Bloom Comb Trees. Designed to support appending new datasets dynamically by updating Comb trees without complete rebuilds, but inherently lacks deletion support.

**SeqOthello** [26] is a hierarchical indexing structure that makes use of Othello hashing. SeqOthello groups similar *k*-mer sets and uses a compression, based on the respective *k*-mer distribution, for each. These *k*-mer distributions are first created for groups of samples, which are then merged. At the root of the hierarchical data structure, a *k*-mer is hashed to its group.

**VariMerge** [27] extends the Vari data structure using a divide-and-conquer approach, splitting large datasets into smaller sets and merging their cdBGs. In addition, VariMerge supports sample insertions.

Despite all this work, it remains a challenge to query the aforementioned public databases. To address this challenge, we previously introduced the Hierarchical Interleaved Bloom Filter (HIBF) [30]. This indexing technique has been implemented in the tool Raptor and has demonstrated significant advancements compared to other commonly used tools such as Bifrost and Mantis by indexing 100 GiB of Ref-Seq data with less than half the storage space, improved query and build times, and comparable or better accuracies. Moreover, the HIBF was capable of indexing one million samples, a feat that has not been achieved before.

One major limitation of the HIBF is its lack of support for dynamic operations: it currently does not support the insertion or removal of sequences or sets without undergoing a complete data structure rebuild. This is a highly resource-intensive operation, which can take hours, especially when dealing with a large number of samples. The necessity for dynamization is evident, as it is desired to stay up to date with the continuously growing databases.

In this work, we propose an involved partial rebuilding technique that facilitates efficient update operations to the HIBF, minimizing additional memory consumption and maintaining a good query time.

## 2 Methods

In the following, we will provide an overview of the Hierarchical Interleaved Bloom Filter (HIBF) [30], including its constituent data structures and its essential operations, building and querying. Next, we will introduce dynamization terminology and objectives that influenced the design decisions (Section 2.2).

In Section 2.3, we will introduce the core principle of the dynamic HIBF (DHIBF): the inclusion of *empty* bins. While the regular HIBF uses all available space to distribute user bins (in the literature also called samples, colors, references, and documents), the DHIBF reserves some space for later use. This reserved space is the *empty* bins and is present in each IBF. To insert a new sample, we first need to determine a suitable insertion location. In the easiest case, we already have enough *empty* bins available. If not, we may resize an IBF or partially rebuild a subtree of the DHIBF. In a second step, we can now insert the sample into the insertion location and validate the false positive rate constraints. In case of a violation, we will rebuild the affected subtree. Figure 4 provides an overview of the procedure.

This will establish a solid groundwork for discussing the individual dynamic operations, namely inserting samples, deleting samples, and modifying samples (Section 2.4, Section 2.7, Section 2.8). Lastly, we will address partial rebuilding, an integral part of the aforementioned operations to ensure good update times (Section 2.5).

### 2.1 HIBF

The HIBF is a *k*-mer based sequencing index. The input sequences for each sample (also referred to as user bins hereafter) are broken down into their representative *k*-mers. We continue to refer to *k*-mers, but minimizers can be read in their place. In brief, Bloom filters (BF) hash *k*-mers of a single sample in a bitvector using a specified number of hash functions. Because different *k*-mer can share hash values, false positives can occur. The false positive rate (FPR) depends on the bitvector’s size, the number of occupied bits, and the number of hash functions [31]. The probability of obtaining a false positive result *p*_*fpr*_ for a single sample depends on the number of stored *k*-mers *n* (*capacity*), the number of hash functions *h* and the bin size *m* (bits) of its Bloom Filter and can be approximated by:

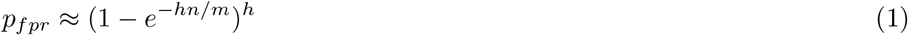

Seiler *et al*. introduced the Interleaved Bloom Filter (IBF) [32, 33]. The IBF contains a BF for each sample, which are of the same length and use the same hash functions, such that their bitvectors can be interleaved (Figure 1). Put differently, each bit in the BF is replaced by a sub-bitvector, with each bit therein corresponding to a different sample. Consequently, the largest sample determines the size of the individual BFs and thereby the overall size of the IBF to guarantee a maximal FPR. When querying a single *k*-mer, the sub-bitvectors at the hashed locations are retrieved and combined using bitwise AND operations. The resulting vector indicates for each sample whether the *k*-mer is present. When querying multiple *k*-mers, these binary vectors are accumulated. For each count, a threshold is applied to decide whether the queried sequence is present or not.

**Figure 1.**
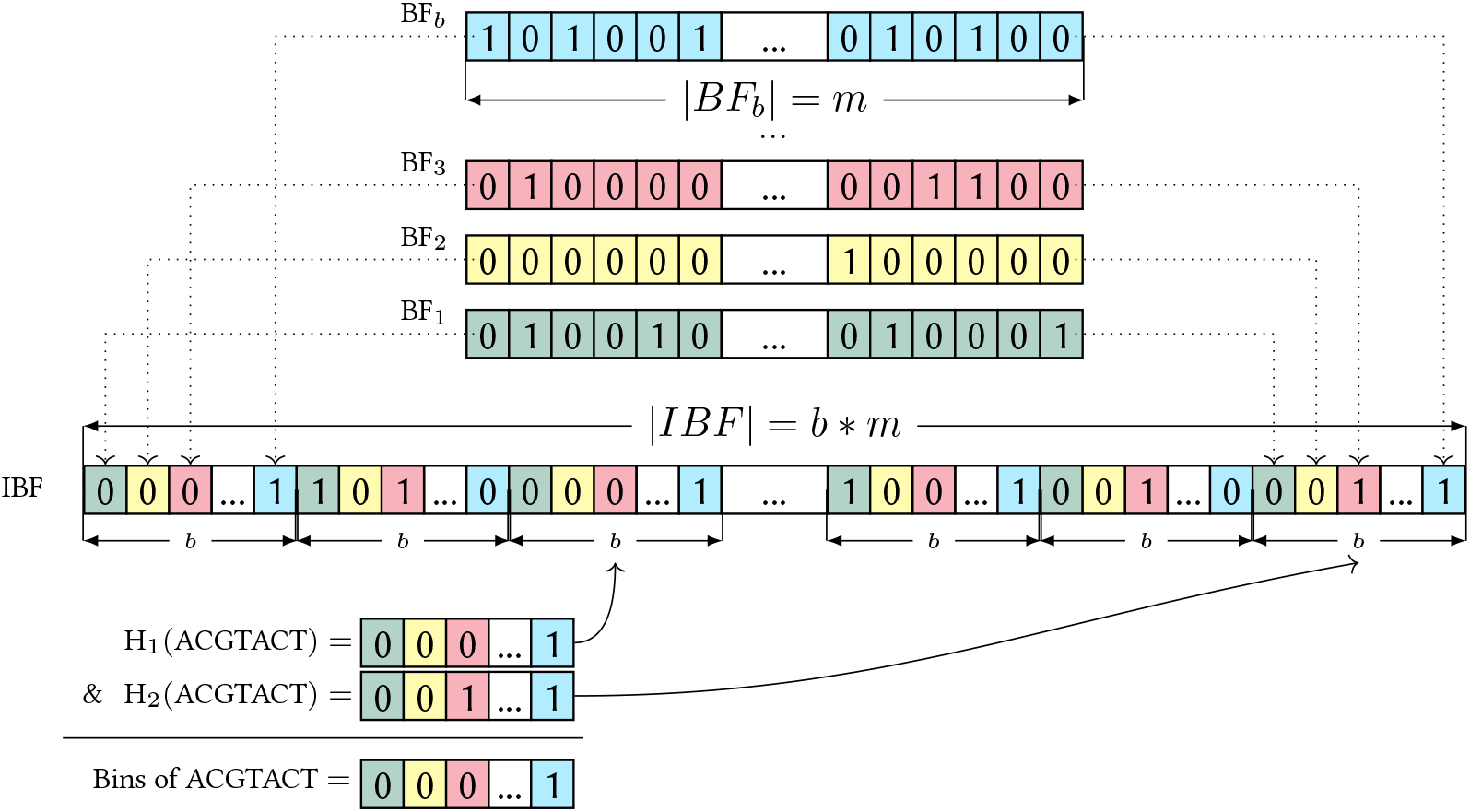
Structuring and querying an Interleaved Bloom Filter (IBF). Differently colored BFs of bin size *m* for *b* samples (shown on the top) are interleaved and yield an IBF of size *b × m*. In the example, *h* = 3 different hash functions are used to store and query a *k*-mer. When inserting a *k*-mer from bin *B*_*i*_ into the IBF, all *h* hash functions are computed, pointing to the positions of the corresponding *h* sub-bitvectors, in which the *i*^th^ bits are set to 1. When querying a *k*-mer, *h* sub-bitvectors are retrieved and combined using a bitwise AND to obtain the binning bitvector of length *b*, where a 1 indicates the presence of the *k*-mer in the corresponding bin. The figure has been adapted from [32].

The IBF faces two limitations. Firstly, the runtime deteriorates when the number of samples becomes large. Think of RNA-Seq collections of over 50,000 samples or contigs in de Bruijn graphs of large genomes, which can easily reach similar numbers [30]. Secondly, the size of the index depends on the maximum sample size in the collection and does not scale well if a high variance in sample sizes exists, for instance, in metagenomic data including different taxonomic domains.

The HIBF addresses these limitations. To improve space consumption and query time, the HIBF uncouples the samples provided by the user, the *user bins*, from the separate bloom filters stored within the IBF, the *technical bins*. This allows for more flexible IBFs with *split bins*, containing partial samples, and *merged bins* containing the content of multiple samples. The HIBF splits large samples over multiple technical bins and merges small samples into a single technical bin. This way, the unevenly distributed sample sizes are evened out in a local IBF, thereby optimizing space consumption. When splitting a sample evenly over *s* technical bins, assuming independent false positive events and an identical *p*_*fpr*_ across all *s* bins, the probability of obtaining a false positive answer for the user bin undergoes a multiple testing correction, resulting in an overall FPR of 1 *−* (1 *− p*_*fpr*_)^*s*^. To distinguish the content of samples that have been merged, each merged bin will be connected to an additional *lower-level* IBF. This IBF needs only to be queried if the merged bin contains the queried *k*-mer, thereby also improving query time for typical queries.

Notably, a HIBF with an optimal space consumption may not have an optimal expected query time and vice versa, representing a complicated interdependent trade-off. It is therefore not straightforward to compute a sensible *layout*, i.e., the organization of samples among a to-be-defined number of technical bins in a collection of IBFs. To solve this, Mehringer *et al*. [30] engineered a *dynamic programming* algorithm that optimizes the space consumption while being constrained by a maximum false positive rate 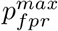, a merged-bin-specific false positive rate 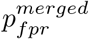, and a fixed maximum number of technical bins *t*_*max*_ for each local IBF. The 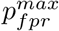 is provided by the user and reflects the desired maximal false positive rate when querying a single *k*-mer. Because false positive results in merged bins do not lead to wrong results but rather to verification in lower-level IBFs, merged bins are allowed to have a higher allowed false positive rate 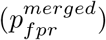. *t*_*max*_ regulates the width of the layout. It restricts the size of each single IBF and thereby ensures that it can be queried efficiently. The dynamic programming algorithm aims to group bins similar in size and, optionally, sequence content in order to minimize the overall size of the HIBF. Placing similar-sized bins in a single IBF avoids wasting space, and merging samples that share *k*-mer results in smaller merged bins. To estimate sizes and sequence similarities of the input data and merged bins, *HyperLogLog* sketches are used [34].

The precomputed layout forms a blueprint for building the actual index (Figure 2). The parallelized *bottom-up* construction algorithm initiates at the top level of the layout and recurses into lower levels until it reaches an IBF with no more merged bins. It then extracts the *k*-mers, computes the actual bin size, and hashes the *k*-mers into the IBF. The union of these *k*-mers is passed on to the merged bin at the next higher-level IBF.

**Figure 2.**
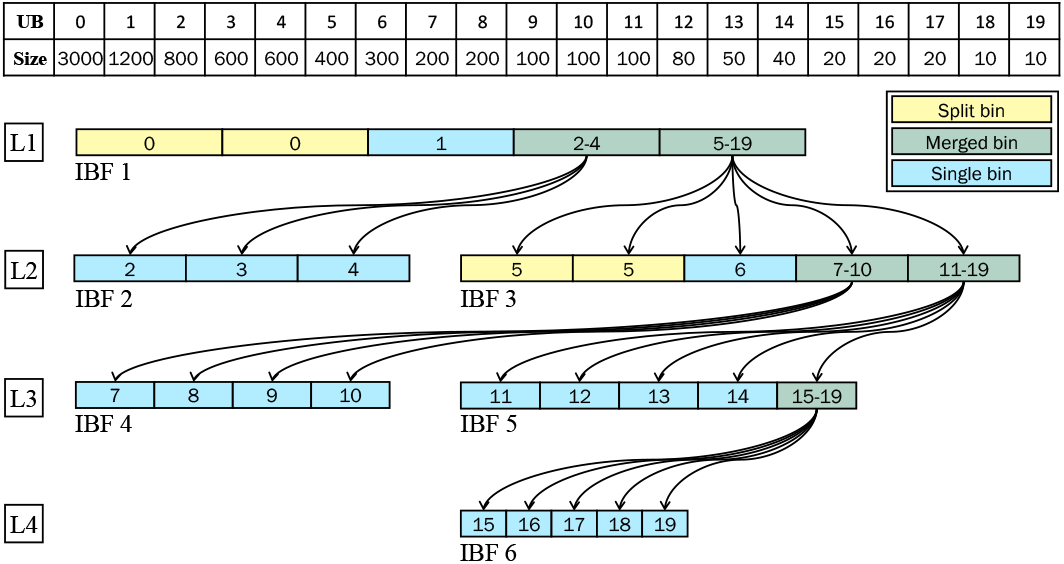
The hierarchical index structure of the HIBF. This exemplary HIBF of 19 numbered user bins (UB) consists of four levels. The top-level (L1) has always a single IBF (IBF-1) with exactly *t*_*max*_ technical bins, in this case 5. This IBF stores the complete dataset using notably less space than a regular IBF because of its split and merged bins (yellow and green, respectively). Each IBF displays boxes representing its technical bins, which store the *k*-mer content of the labelled user bins. For instance, in IBF-1, UB-1 is split and stored in two technical bins, UB-2 in a single technical bin, and UB-2 to 4 are merged into one technical bin. Note that the schematic does not depict that in reality these technical bins are interleaved. IBFs in lower levels each correspond to a merged bin in a previous level. For instance, IBF-2 and IBF-3 on the second level (L2) correspond to the first and second merged bin of IBF-1. Note that the IBFs in the layout form a tree, where the root is the top-level IBF and the leaves consist of IBFs without merged bins. The figure was adapted from [30].

### 2.2 Dynamization objectives

In this section, we will touch upon the complexity of incorporating dynamic operations in the HIBF and the various trade-offs that have to be kept in mind. We will introduce terminology that will aid the description of the design choices made throughout the various update operations discussed afterward.

Efficient support for dynamic operations should avoid frequent full rebuilds of the index. However, allowing for reasonable update times generally comes at the cost of increased space usage or query time. The computation of an optimal layout for the HIBF can be conceptualized as a constrained optimization problem. Its objective function would capture the trade-off between query time and space consumption. In the case of the dynamic HIBF, the objective becomes even more complex due to the consideration of (cumulative) update time. Moreover, these objectives are interdependent, scale differently with the index’s size, and should contribute differently to the objective function depending on the application. For instance, applications may differ in the available computational resources and relative frequencies of updates and queries. Lastly, this optimization problem should be solved efficiently to allow for fast index construction.

We will describe the anticipated impact of the proposed techniques and assess their effectiveness for various use cases. In the following, we will use the terms *feasible* and *relaxed* to refer to different configurations of the HIBF, representing various aspects of the optimization problem. Firstly, a state is considered *feasible* if it meets the constraint that the actual false positive rate *p*_*fpr*_ for each user bin is below the threshold, i.e. 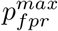 for split bins and 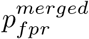 for merged bins.

Secondly, the index is in a *relaxed* state if it can tolerate a certain number of update operations without requiring a costly rebuild. This can, for instance, be achieved by allocating additional space to accommodate fast insertions.

### 2.3 Empty Bins

*Empty bins* are technical bins that are not yet associated with any sample, and the corresponding Bloom filters are thus empty. These empty bins can accommodate new samples. When the index has sufficient empty bins to perform a certain number of updates, it is in a relaxed state. Having empty bins in the layout is therefore an integral part of the dynamization method (Figure 3). We adapted the dynamic programming layout algorithm from [30] to reserve a user-given percentage, *ϕ*_*∅*_ *∈* [0, 1), of *t*_*max*_ to be used as empty bins. While the individual IBFs are still constructed with *t*_*max*_ technical bins, the layout is computed with *t*_*max*_ *\** (1 *− ϕ*_*∅*_) technical bins. This approach achieves robust runtime guarantees, relying less on the data being used.

**Figure 3.**
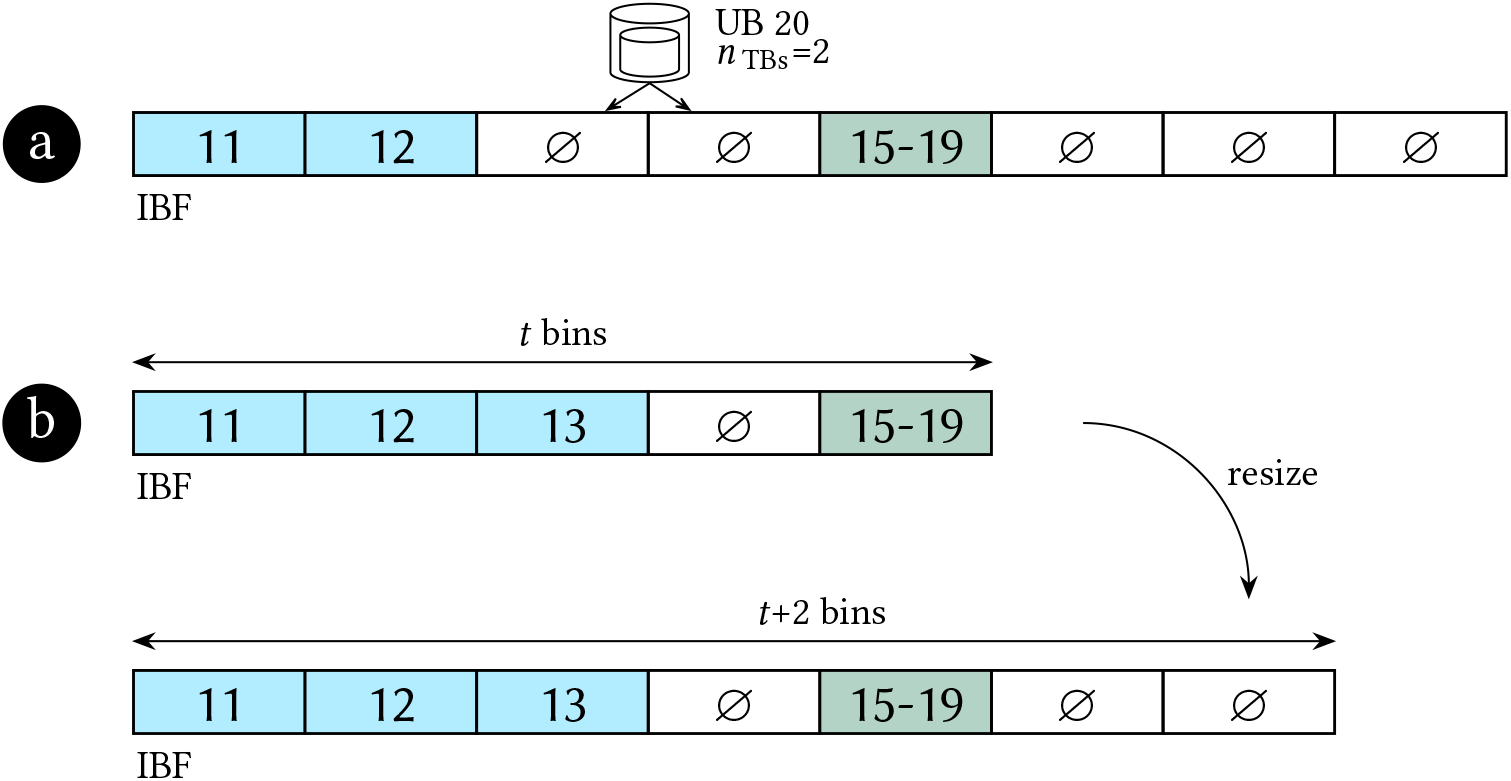
Selecting IBFs for sample insertions. Empty bins are denoted by ∅. After selecting an IBF, the algorithm scans an auxiliary table from left to right to find adjacent empty bins within the IBF. In example **(a)**, there are enough (*≥ n*_TB_) adjacent empty bins to accommodate the new sample. In case insufficient adjacent empty bins are available (**b**), a resize operation is executed. Enough empty bins (EB) are added such that the new sample can be accommodated. However, the number of technical bins is not allowed to exceed *t*_*max*_ + 64.

### 2.4 Inserting samples

Inserting a new sample consists of two major steps. Firstly, a suitable insertion location must be found (Section 2.4.1). Afterwards, the sample can be inserted at the found location, potentially triggering a rebuild (Section 2.4.2). A high-level overview of the proposed dynamic insertion operations is depicted in Figure 4.

**Figure 4.**
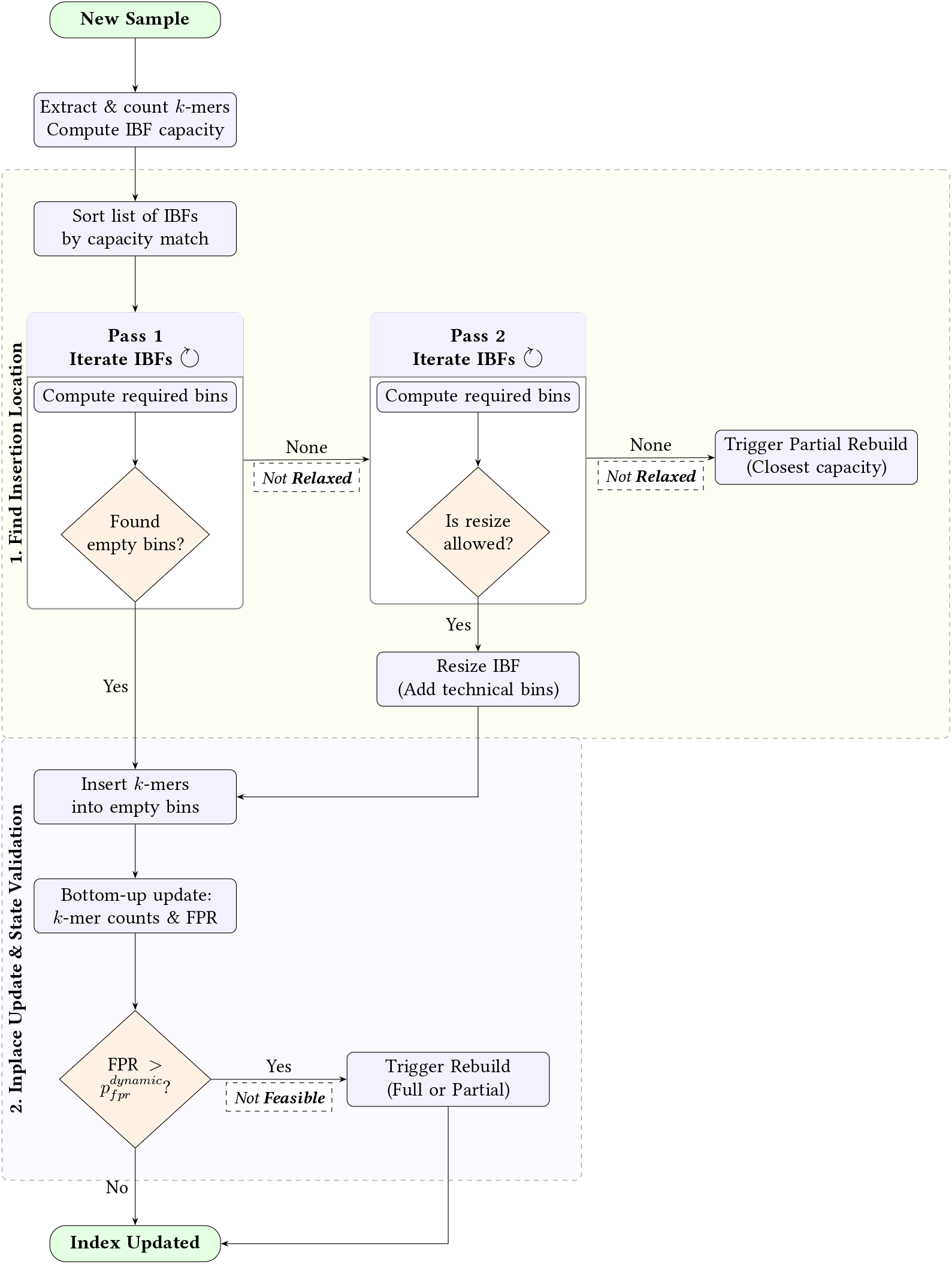
Schematic overview of the dynamic HIBF sample insertion. The process ensures the index remains in a *relaxed* and *feasible* state by searching for adjacent empty bins, opportunistically resizing IBFs to prevent immediate bottlenecks, and evaluating dynamic FPR thresholds 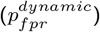 to trigger necessary rebuilds.

When a new sample is to be inserted in the index, its *k*-mers are extracted and counted. Then, we compute the capacity *n* of each IBF, i.e., the maximum number of elements each IBF can accommodate while still maintaining the FPR constraints (Equation 2).

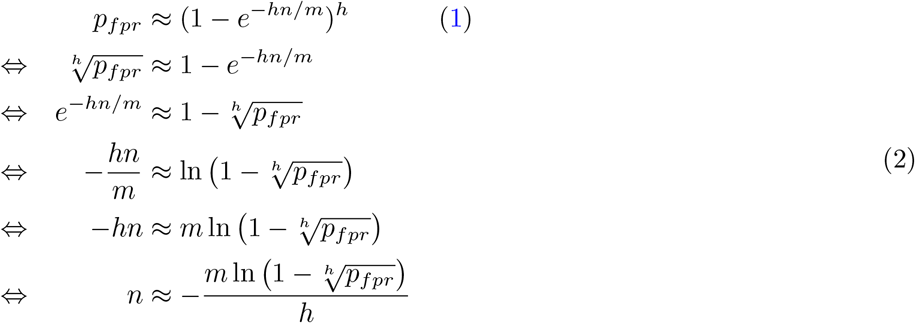

#### 2.4.1 Finding the Insertion Location

Next, we search for an IBF to insert the new sample into. We sort the list of IBFs by how closely their capacity matches the *k*-mer count and evaluate each IBF until we find a suitable location. The evaluation begins by computing the number of technical bins needed to store the new sample. If the current IBF has sufficient consecutive empty bins available, it is selected as the insertion location (Figure 5a). In case no IBF is suitable, we repeat the evaluation process with the added possibility to resize an IBF (Figure 5b). Resizing an IBF allows us to add more technical bins to an existing IBF without constructing it from scratch. This is much faster than (partially) rebuilding the full index, albeit only when done sparsely. We allow resizing up to *t*_*max*_ + 64 technical bins. Resizing is not allowed in three cases: 1) the IBF has already been resized, 2) the IBF is the root IBF, 3) the IBF has been part of a partial rebuild. If this second search is unsuccessful, a partial rebuild for the IBF with the closest capacity is triggered. The partial rebuild will make the IBF and its children ineligible for future resizing.

**Figure 5.**
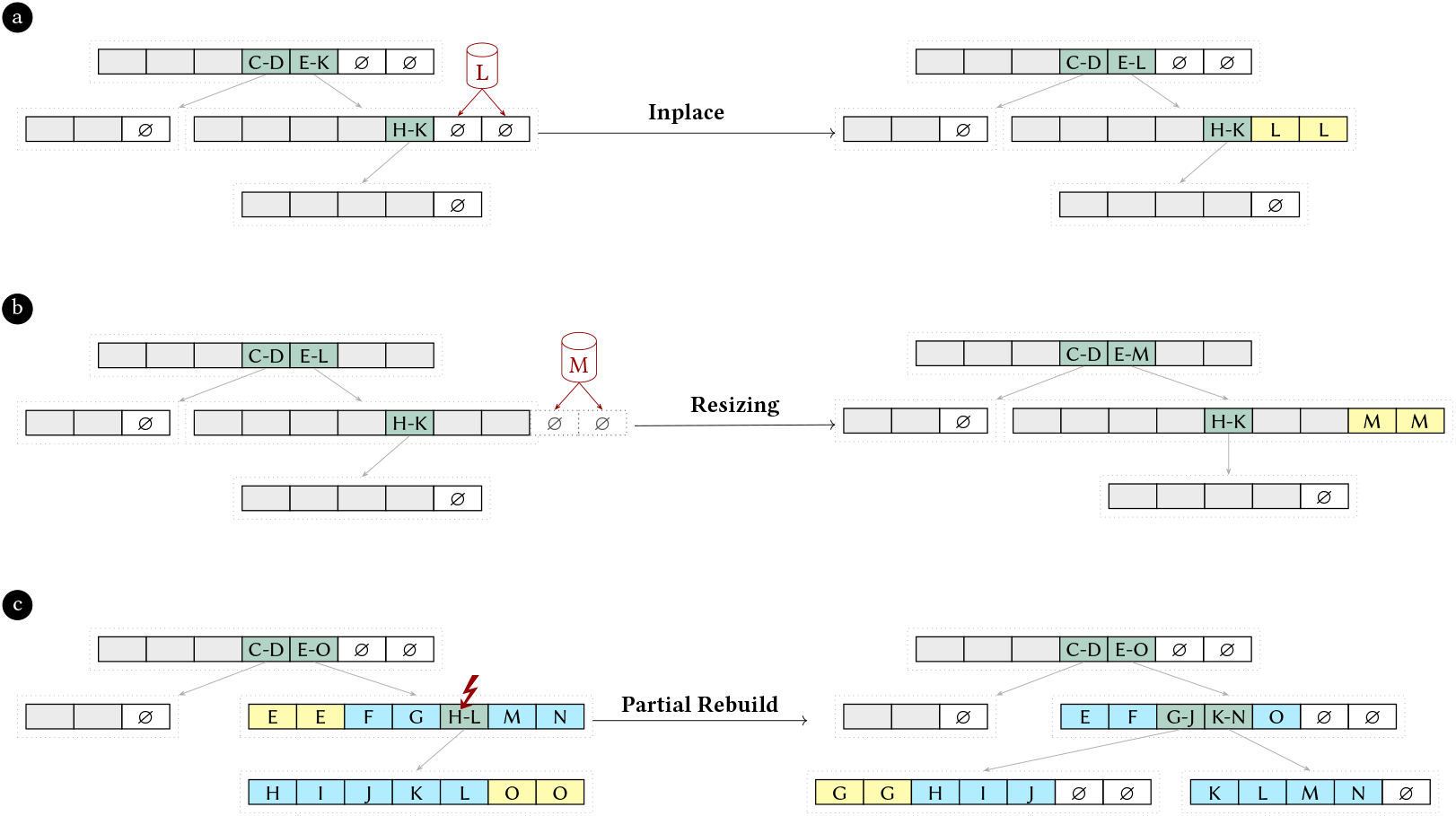
Sample insertion examples. Empty bins are denoted by ∅. A gray background is used for present bins that are not relevant for the respective example. A merged bin exceeding the false positive threshold after sample insertion is marked with a f. The left-hand side shows the initial index status, while the right-hand side shows the index after insertion. a) depicts an inplace insertion of user bin *L*. b) shows how the user bin *M* is inserted by resizing an IBF. c) illustrates a partial rebuild triggered by exceeding 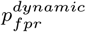 at the merged bin *H − L*.

#### 2.1.2 Inplace Update

The *k*-mers are inserted into the empty bins while updating associated auxiliary metadata. If the target IBF was not the root IBF, this process is repeated for the parent IBF until we arrive at the root IBF.

While inserting *k*-mers, we also track the effective number of elements added to each technical bin to monitor its false positive rate (FPR). Specifically, we only increment the bin’s element counter if inserting the *k*-mer sets at least one previously unset bit, i.e., the incoming *k*-mer is not reported as present by the Bloom filter. Because a Bloom filter cannot distinguish between a duplicate *k*-mer and a false-positive hash collision of a novel *k*-mer, this counter represents the number of bit-modifying insertions rather than the exact number of distinct *k*-mers. However, because *k*-mers that already test positive do not set any additional bits, they do not increase the bitvector’s saturation or degrade the actual FPR. Substituting this effective element count *n* into Equation 1 therefore provides a close estimate of the technical bin’s current FPR, particularly for merged bins where samples share *k*-mers, without overcounting redundant insertions. If the estimated FPR exceeds the allowed threshold for any IBF during insertion, we record the IBF’s location to trigger a partial rebuild once the insertion completes. The FPR threshold depends on the type of technical bin the insertion occurs in and, for merged bins, on its level (depth within the tree). Split bins have to conform to 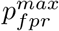. For merged bins, we use the merged-bin-specific FPR 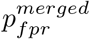 with a level-dependent relaxation factor:

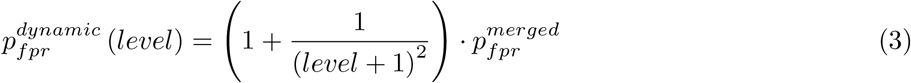

The *level* is 0 for the root IBF and increases by one for each merged bin encountered along the path to an IBF. The dynamic FPR 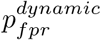 accounts for the presence of more children on higher levels while not precluding partial rebuilds of lower-level subtrees. Note that some form of relaxation factor is necessary for dynamization. When an HIBF is constructed, it is likely that the fullest bin is a merged bin, in which case it 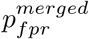 is used to determine the required bin size. As this is a strict lower bound for the bin size, any insertion into the merged bin would immediately trigger a rebuild.

After selecting an IBF, the algorithm looks for empty bins within this IBF using an auxiliary table (Figure 3, Figure 5a). If there are not enough adjacent empty bins available, a resize and possibly a rebuild operation (Figure 5c) will be triggered.

Finally, the *k*-mers are hashed into the empty bins, and auxiliary tables with FPRs and *k*-mer counts are updated for the bins along the insertion path. The insertion into the hierarchical index is done bottom-up and the auxiliary tables are updated simultaneously. To update the *k*-mer counts of the merged bins, we check for each new *k*-mer if it was already present when inserting it. Only the number of *k*-mers that were not already present is added to the count. If any of the FPRs exceed the FPR threshold, a rebuild operation is triggered (Figure 5c).

#### 2.4.3 Resizing IBFs

When resizing the IBF, the number of technical bins is increased. If there are no empty bins available or, more generally, the requested number of technical bins is greater than the potential technical bins already allocated for, a reallocation is required. The reallocation is effectively a backwards-copy, efficiently shifting every interleaved block and setting the extra bits to zero. When reallocating, we increase the requested number of technical bins to the next multiple of the word size. All auxiliary tables are resized accordingly. See Figure 3b for an example.

A resize operation is triggered when a new sample needs to be inserted into an IBF that does not have sufficient adjacent empty bins for inserting new samples. In theory, resizing might be considered redundant because the rebuild operation, explained in the next section, restores the empty bins.

However, in practice, resizing is still beneficial because samples share sequence content and are smaller than what their technical bin can accommodate, so consequently, their merged bins will also not have reached the FPR threshold. In addition, the child IBFs may not grow equally, making regular rebuilds of the subtree inefficient. Furthermore, there may be situations where multiple adjacent empty bins are required for inserting a new sample, while only scattered empty bins exist due to previous deletions.

### 2.5 Rebuilding

Here we describe the different rebuilding operations that are triggered within the update operations when the index grows out of balance, and the FPR threshold or *t*_*max*_ is exceeded. These rebuilding operations should restore a relaxed state.

If the top-level IBF is resized, a complete rebuild of the index is necessary. Otherwise, the hierarchical structure will grow more in width than in depth, and query times and space usage will diverge from the balanced state. Therefore, a complete rebuild is triggered if the number of bins at the top level exceeds *t*_*max*_ or any bin exceeds the FPR threshold after insertion (see [30] for further details). Similarly, resizing may also prompt a partial rebuild operation on lower levels when *t*_*max*_ or the FPR threshold is exceeded.

A full rebuild may also be triggered if the number of resized or partially rebuilt IBFs exceeds twice the number of IBFs of the original, built-from-scratch HIBF. This acts as a stopgap to prevent excessive partial rebuilds.

A full rebuild constitutes a rebuild of the index from scratch. For partial rebuilds, we repurposed the dynamic programming, rearrangement, and hierarchical build algorithm from [30] for (partially) rebuilding with empty bins. These algorithms require the relaxed state of a rebuilt (sub)tree to accommodate a fixed minimum percentage of additional samples.

### 2.6 Batch insertions

When inserting multiple samples at once, we iteratively insert one sample. However, if at any point a full rebuild is triggered, we include all to-be-inserted samples in this full rebuild.

### 2.7 Deleting samples

When a sample is deleted, the corresponding technical bins are cleared, and the auxiliary tables are updated accordingly. Note that its *k*-mers stored in its parent merged bins are only removed during later rebuild operations. If these merged bins have not reached the FPR threshold, new samples can now be inserted in the freed space.

### 2.8 Modifying samples

Deleting a sample requires only emptying all its (lowest-level) technical bin(s). Afterward, any modified sequence content could be inserted in the same location. Bloom filters do not support deletions of individual *k*-mers, as it is impossible to determine if a set bit originated from the *k*-mer to be deleted. Therefore, any of its parent merged bins will not be modified, but their excess *k*-mers will be deleted later during a rebuild operation.

## 3 Results

The dynamic version of the Hierarchical Interleaved Bloom Filter has been integrated into the Raptor tool [32], enabling its use with the static HIBF and the dynamic HIBF. The latter integration allows for efficient, incremental updates to the underlying data structure, making it well-suited for large-scale, fast, ever-growing sequence datasets like NCBI SRA, GenBank, and RefSeq, among others. In this section, Raptor-Static refers to the original HIBF implementation, and Raptor-Dynamic refers to the dynamic HIBF.

To evaluate the dynamic capabilities of the HIBF, we first conducted experiments using two simulated sequence datasets, each containing 100,000 sequences and totaling 1 Gbp of data (see Appendix Figure 13). The first dataset features an even distribution of random DNA sequences, each 10,000 bp in length. The second dataset emulates a RefSeq-like sequence-length distribution, derived from the sequence lengths in RefSeq [9]^[1]^ by sampling and normalizing for a 1 Gbp total size^[2]^.

For the evenly distributed and RefSeq-like simulated datasets, we generated 100,000 and 97,146 reads, respectively, of 250 bp each with two substitution errors. The number of reads simulated per sequence is proportional to the sequence length. In addition to synthetic data, we evaluated performance on real-world human RNA sequencing data, comprising 39,400 samples and totaling 110.6 TB of compressed data. This dataset^[3]^ was originally introduced in the dynamic Mantis study [29]. We further evaluated the indexing capabilities of the methods based on Bacterial and Archaeal genomes from RefSeq.

We compared the dynamic HIBF implementation in Raptor-Dynamic configured with 10% empty extra bins against several state-of-the-art tools: the non-dynamic HIBF (Raptor-Static), Metagraph [24], COBS [17], and Fulgor [18], as well as the dynamic methods kmindex [20] and Bifrost [16]. Those methods were chosen based on their ability to index large genomic data sets efficiently and with an active code base and user support. Although Metagraph offers a dynamic mode to update indices, the authors discourage its use due to instability and performance issues^[4]^; thus, we used the static version for fair comparison. We attempted to include the dynamic versions of PAC [25] and Mantis [29] in our evaluation, but both tools exhibited critical failures or incomplete functionality during execution: an updated PAC index could not be queried^[5]^, and mantis mst as well as non-bulk queries crashed^[6]^. Despite attempts to contact the respective authors, we were unable to resolve these issues or obtain working implementations in time for publication of this article. kmindex uses minimizers (size 10 by default) to partition the k-mers, not to winnow them, and needs a manually chosen Bloom filter size (set to 30,000 bits). Each update builds a new sub-index, and all sub-indices are queried together, because merging them failed^[7]^. As kmindex always indexes all k-mers, we evaluated it on the simulated data only, where Raptor does the same (it uses minimizers with a window size of 24 on real data). On real data, Bifrost crashed after at most two updates^[8]^.

Our evaluation of the Raptor-Dynamic follows an incremental update paradigm: we begin with an initial set of sequences and progressively update the index with new batches of sequences. For Raptor-Static, Metagraph, COBS, and Fulgor, each update was performed from scratch by re-indexing all sequences up to the current batch. Raptor-Dynamic, kmindex, and Bifrost instead update their index. The Run Number in the figures refers to the update batch, with 0 being the initial build. All tools used 24 threads and a k-mer size of 20.

Figure 6 presents the build plus query time and memory usage for the simulated even dataset; Table 2 reports build and query times separately. Note that results are presented in a logarithmic scale for each iteration of incremental updates/rebuilds. There are substantial differences in scaling behavior among the evaluated tools. Bifrost, COBS, Fulgor, Metagraph and Raptor-Static exhibit a higher increase in time with each update, indicating limited efficiency in handling incremental data additions, with higher processing time as the index grows. The build time of kmindex stays constant, but its query time grows. Raptor-Dynamic, leveraging the dynamic HIBF, consistently requires very little time and memory to update and query the index, maintaining an average low build time throughout the experiment. There are occasional full rebuilds necessary that take time comparable to the rebuild of the other methods. In this scenario, considering all updates/rebuilds, Raptor-Dynamic was 6.5 times faster to build and query than COBS, the second fastest method (average of 18 seconds and 117 seconds, respectively), and 11.5 and 49 times faster than the dynamic methods kmindex and Bifrost. Raptor-Dynamic also requires less memory for the incremental updates of each iteration (Figure 6b), requiring, on average, 6.2 times less memory than COBS. COBS achieved the smallest index sizes, followed by both Raptor variants and Bifrost. Raptor-Dynamic needs slightly larger indices than Raptor-Static to accommodate empty bins for the dynamic updates (on average 3.6%, at most 2.2 times after the full rebuild in the last run, Figure 6c). kmindex produced by far the largest index (589 GiB).

**Table 2.**
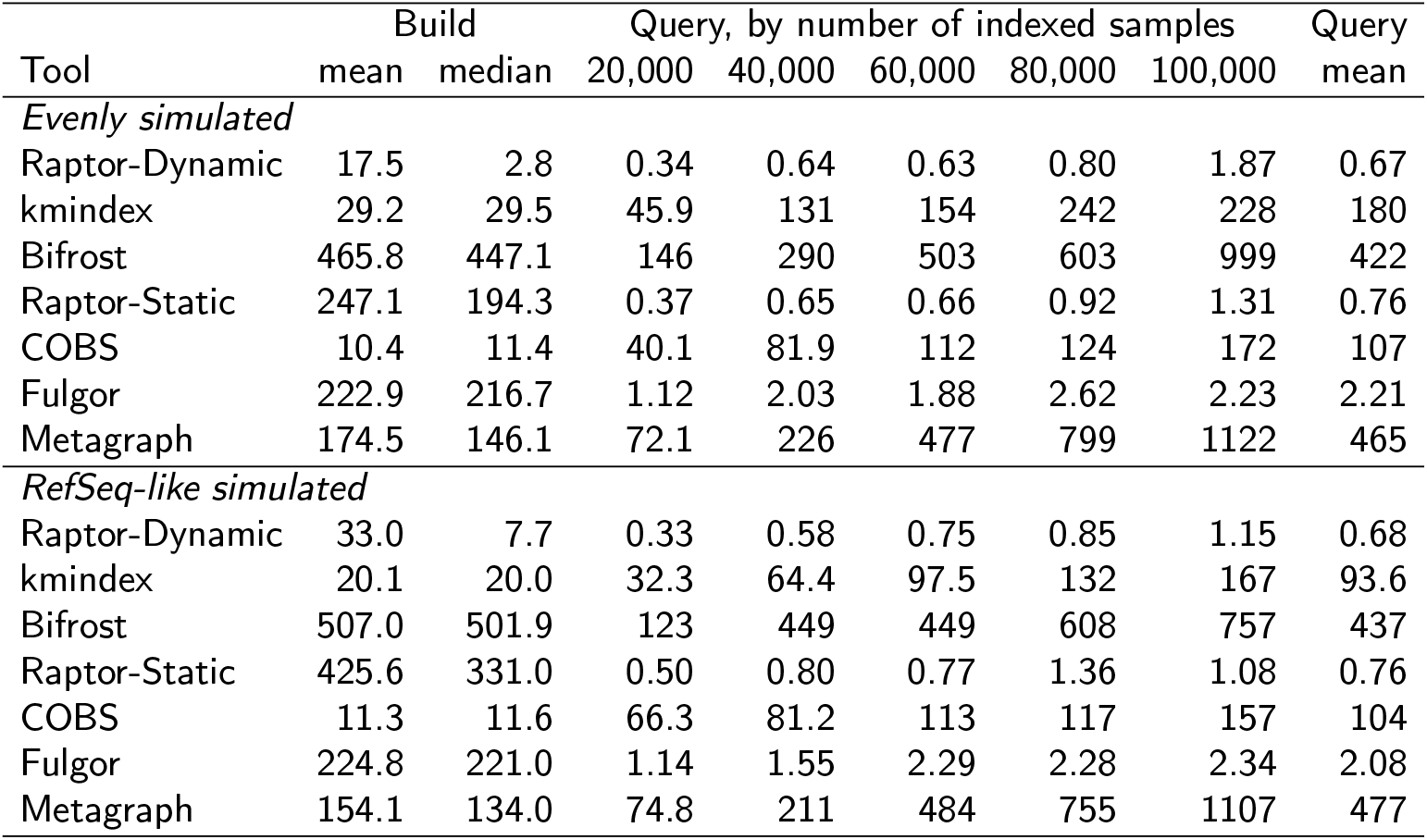
Build and query times on the simulated datasets. Times in seconds over runs 1–90: build time per update and time to query all reads at the given number of indexed samples (runs 10, 30, 50, 70, and 90). Tools updating an index are listed first.

**Table 3.**
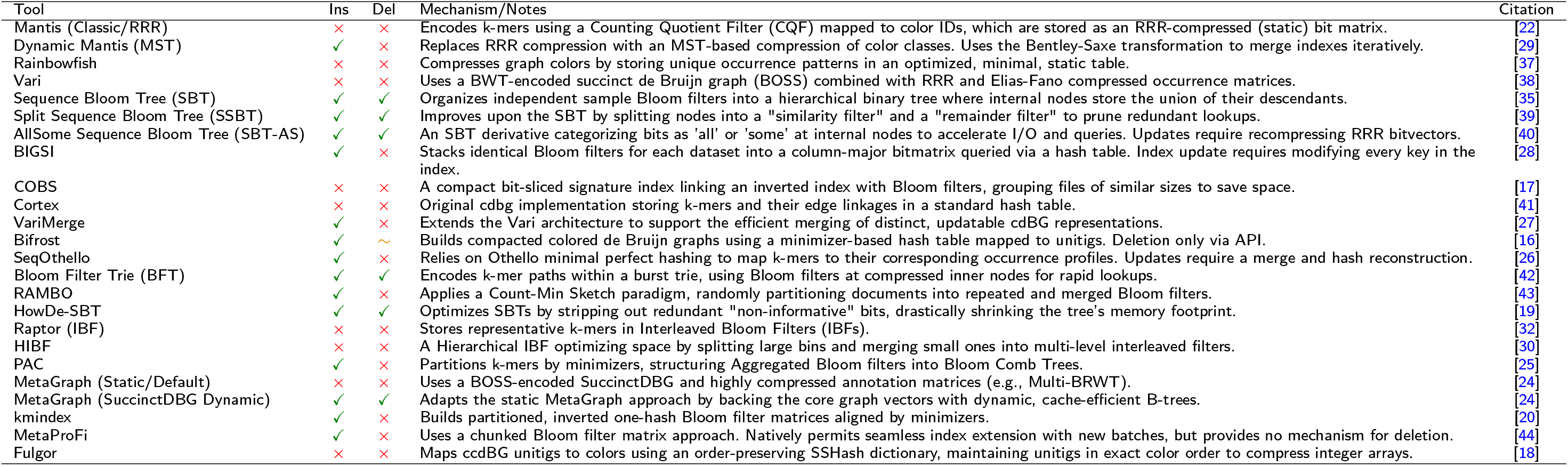
Algorithmic comparison of multi-set sequence indexing tools. Overview of representative colored de Bruijn graph and sequence aggregation indices, detailing their support for dynamic insertion and deletion operations along with their core indexing mechanisms. Expanded version of Table 1.

**Figure 6.**
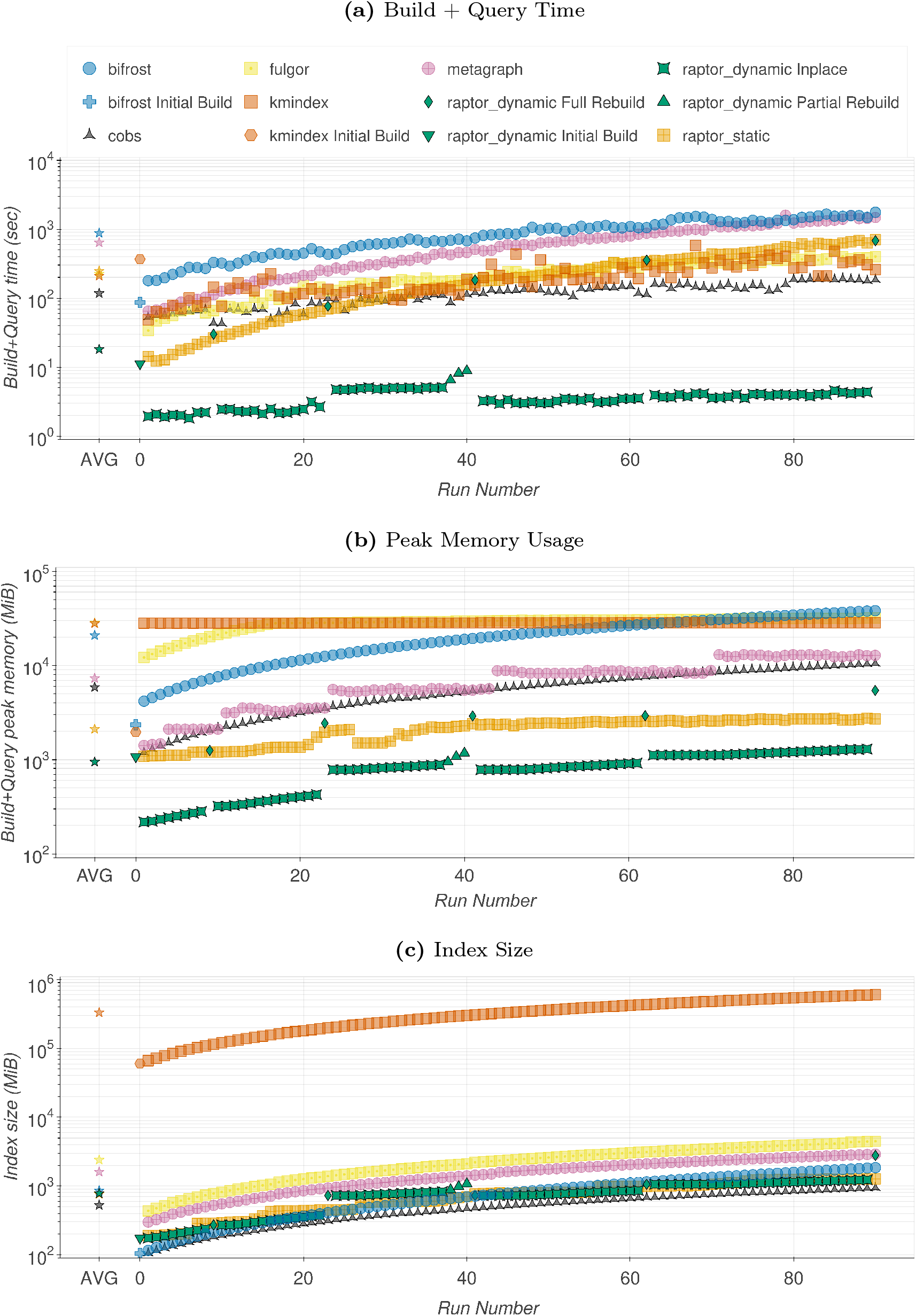
Batch Update of Evenly Simulated Dataset. Build + Query time in seconds (a), peak memory usage in MiB (b), index size in MiB (c). The initial build (Run Number x=0; Raptor-Dynamic, kmindex, and Bifrost only) contains 10,000 sequences, and each incremental run (x*≥*1) adds 1,000 sequences. Leftmost points are the average among all data points for each tool, color-based.

To further assess the dynamic HIBF’s performance characteristics, we repeated the evaluation using a RefSeq-like dataset designed to mimic the distribution of sequence lengths found in genomic databases. Figure 7 presents the results plotted on a logarithmic scale. The trends observed with the simulated even distribution (Figure 6) are largely maintained with the RefSeq-like data: Raptor-Dynamic was 3.4 times faster to build and query than both COBS and kmindex.

**Figure 7.**
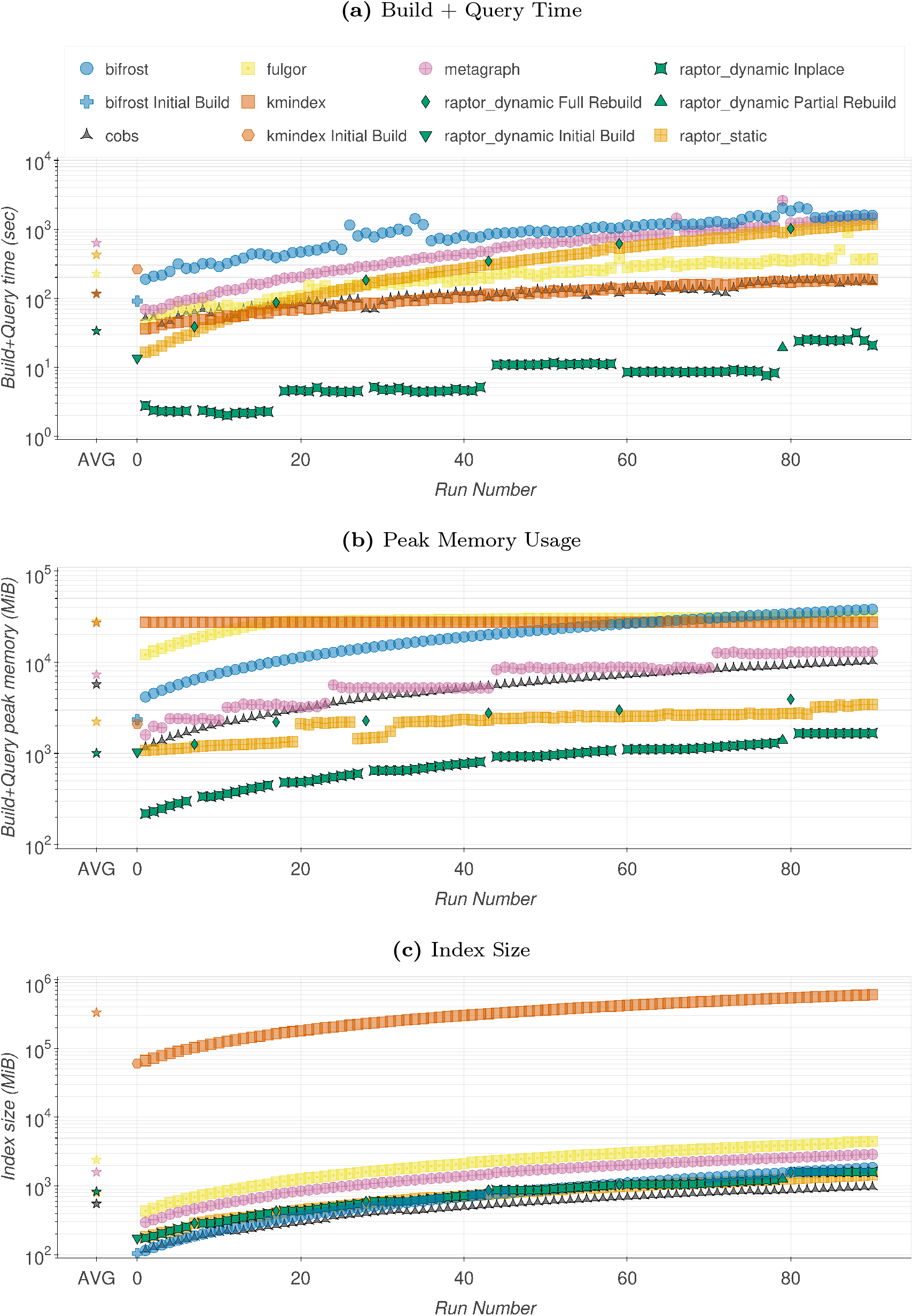
Batch Update of RefSeq-like Simulated Dataset. Build + Query time in seconds (a), peak memory usage in MiB (b), index size in MiB (c). The initial build (Run Number x=0; Raptor-Dynamic, kmindex, and Bifrost only) contains 10,000 sequences, and each incremental run (x*≥*1) adds 1,000 sequences. Leftmost points are the average among all data points for each tool, color-based.

To evaluate accuracy, all simulated reads (100,000 for the even and 97,146 for the RefSeq-like dataset) are queried after every update, with a read’s sample of origin as ground truth. A hit in this sample is a true positive (TP), a hit in any other sample is a false positive (FP). A read without hits is a false negative (FN) if its sample is indexed and a true negative (TN) otherwise, so reads from samples not yet indexed are negative queries (89,000 and 86,453 after the first update, 0 after the last). Raptor requires 191 of a read’s 231 k-mers (k-mer lemma for two errors), and COBS, Metagraph, kmindex, and Bifrost require 80% of a query’s k-mers to match. Fulgor ran in its default mode, since its threshold option produced spurious matches.

When evaluating the results of querying the simulated reads for both even and RefSeq datasets, Raptor-Dynamic, Raptor-Static, COBS, Metagraph, and Bifrost achieve perfect accuracy in all runs, with identical results for both Raptor variants. kmindex only has up to 9 false positives per run on the RefSeq-like dataset (0.003% overall). Fulgor consistently had a significant number of false positives and false negatives, around 4.8% and 1.8% on average for all batches, respectively. The number of false positives gradually declines until it reaches 0 after the last batch, when all samples are in the index.

Queries on an incrementally updated index are on average not slower than on a freshly rebuilt index with identical samples for both Raptor runs (Table 2), and the other tools took 3 to 700 times as long. In-place updates took on average 2.9 and 8.0 seconds on the even and RefSeq-like dataset, but rebuilds account for 85% and 78% of the total build time of Raptor-Dynamic. Its mean build time therefore exceeds that of COBS on both datasets and that of kmindex on the RefSeq-like dataset, although their queries take more than 100 times as long.

Repeating the even experiment with batch sizes of 500, 2,000, and 3,000 samples (Figure 8) shows little effect of the batch size. Full rebuilds were triggered at similar numbers of indexed samples, slightly later for larger batches, since a rebuild includes all remaining samples of the batch (Section 2.6). Only batch sizes of 500 and 1,000 triggered a fifth rebuild near 100,000 samples, which accounts for most of the difference in total build time and final index size. The cost of in-place updates per sample and query times show no clear dependence on the batch size. However, timings vary between runs: the runs with a batch size of 1,000 in Figure 8 and Figure 6 made identical update decisions but differed by a factor of 1.7 in total build time.

**Figure 8.**
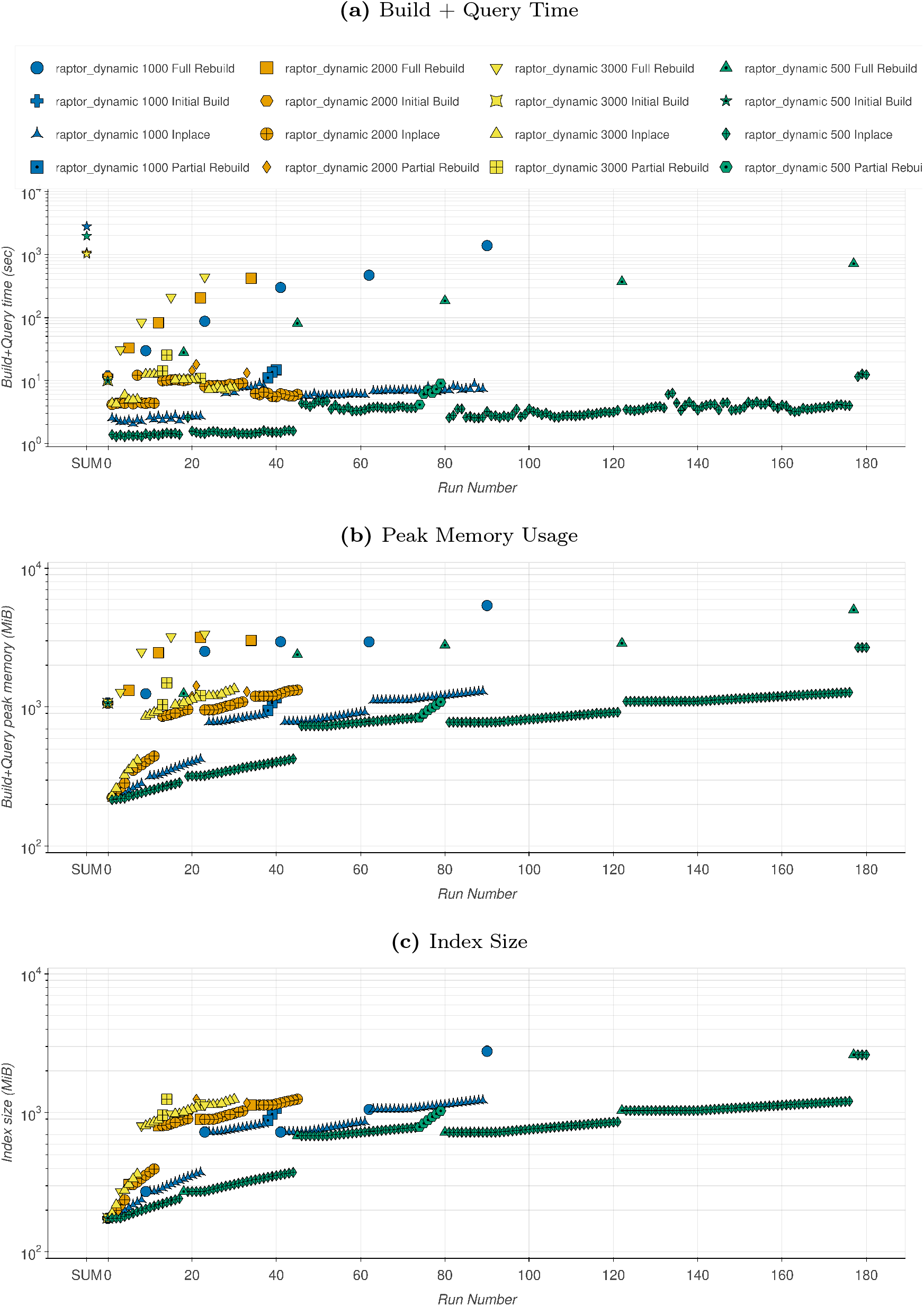
Batch Update of Evenly Simulated Dataset with Different Batch Sizes. Build + Query time in seconds (a), build peak memory usage in MiB (b), index size in MiB (c). The initial build (Run Number x=0) contains 10,000 sequences, and each incremental run (x*≥*1) adds 500, 1,000, 2,000, or 3,000 sequences. Leftmost points are the sum among all data points for each variation, color-based.

For a mixed workload, each run of the even experiment first deletes 100 previously inserted samples and then inserts 1,000 new ones (Figure 9). As a modification is a deletion followed by an insertion (Section 2.8), this exercises all update operations. Accuracy was unaffected: the reads no longer found matched the deleted samples (100 per run), and no false positives occurred. An update took 4.9 seconds in the median. Compared to insertions only, the index was smaller on average (713 and 786 MiB), and queries took 0.84 instead of 0.67 seconds.

**Figure 9.**
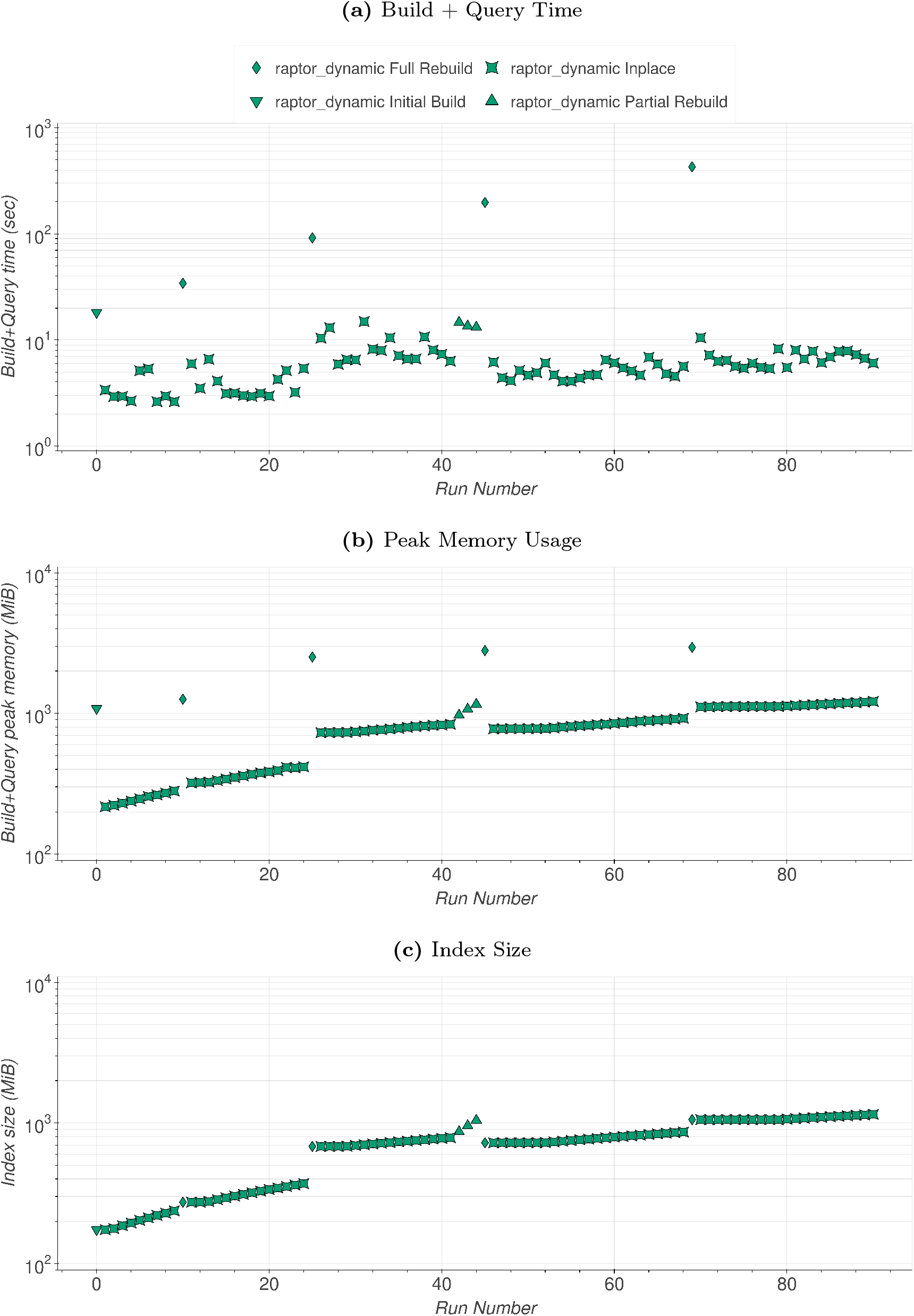
Batch Update with Insertions and Deletions of Evenly Simulated Dataset. Build + Query time in seconds (a), peak memory usage in MiB (b), index size in MiB (c). The initial build (Run Number x=0) contains 10,000 sequences, and each incremental run (x*≥*1) deletes 100 sequences and adds 1,000 sequences.

Next, the dynamic HIBF was evaluated for real data, indexing 39,400 human RNA samples from SRA (Figure 10). To compare Raptor in a timely manner with other methods, we randomly sub-sampled each read file to 1% of its reads, ending up with approximately 1 TiB of compressed data. Note that only build times are reported in this evaluation to enable a more accurate comparison of the methods’ outcomes, since query times steadily increase for each tool in every iteration due to the repetitive nature of human RNA data. Following the same trend of the simulated data analysis, dynamic updates of real-data indices with Raptor-Dynamic provide a clear reduction in time and memory consumption. Raptor-Dynamic required, on average, only 12 minutes to update an index, adding 100 samples on each iteration, while Raptor-Static needed 81 minutes. Accumulating the build times, Raptor-Dynamic took a little more than 80 hours to fully update the index after 385 iterations, while Raptor-Static needed 520 hours.

**Figure 10.**
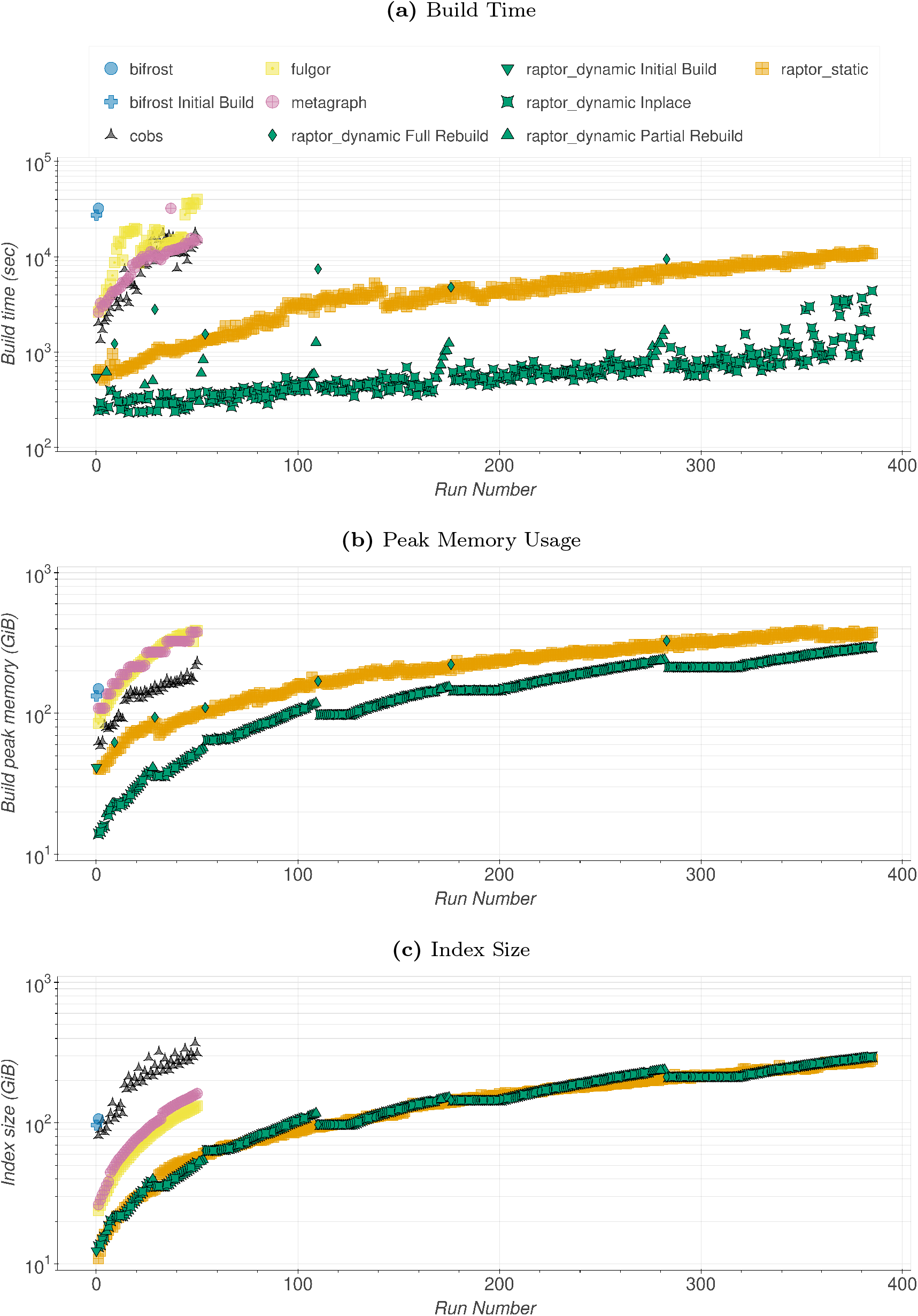
Human RNA sub-sampled to 1%. Build time in seconds (a), peak memory usage in GiB (b), and index size in GiB (c). The initial build (Run Number x=0; Raptor-Dynamic and Bifrost only) contains 1,000 sequences, and each incremental run (x*≥*1) adds 100 sequences. COBS, Fulgor, and Metagraph were executed only for the first 50 batches due to long run times; Bifrost failed after the first update.

We did not further execute Fulgor, Metagraph, and COBS after the 50th iteration due to prohibitive runtimes. Inspecting only the first 50 iterations, Raptor-Dynamic needed, on average, 6.3 minutes for each update, while Raptor-Static took 14.7 minutes, COBS 147 minutes, Metagraph 155 minutes, and Fulgor 264 minutes. In total, Raptor-Dynamic took 5.3 hours to update the first 50 runs, while COBS, the second-best alternative method, took more than 122 hours (approximately 5 days) to perform the same task. With raw human reads data, Raptor indices (static and dynamic) achieved the best compression levels and are the smallest in this evaluation (Figure 10c). During incremental updates, the dynamic HIBF’s peak memory footprint in RAM is roughly equivalent to the size of the existing index, except during occasional full rebuilds, which happen only 6 times out of 385 in this evaluation (Figure 10b).

We further evaluated Raptor-Dynamic’s performance by indexing the full human RNA data without sub-sampling. Results for all 39,400 samples totaling 100 TB of compressed data are presented in Figure 11. Here, Raptor-Dynamic was executed with --use-filesize-dependent-cutoff to remove low-frequency k-mers, using the file-size-dependent thresholds introduced by Solomon and Kingsford [35]. Raptor-Dynamic needed, on average, including the initial build of 1000 samples, 75 minutes to update the main index, inserting in consecutive batches of 100. The final index with all queryable 39,400 human RNA samples is 394 GiB in size.

**Figure 11.**
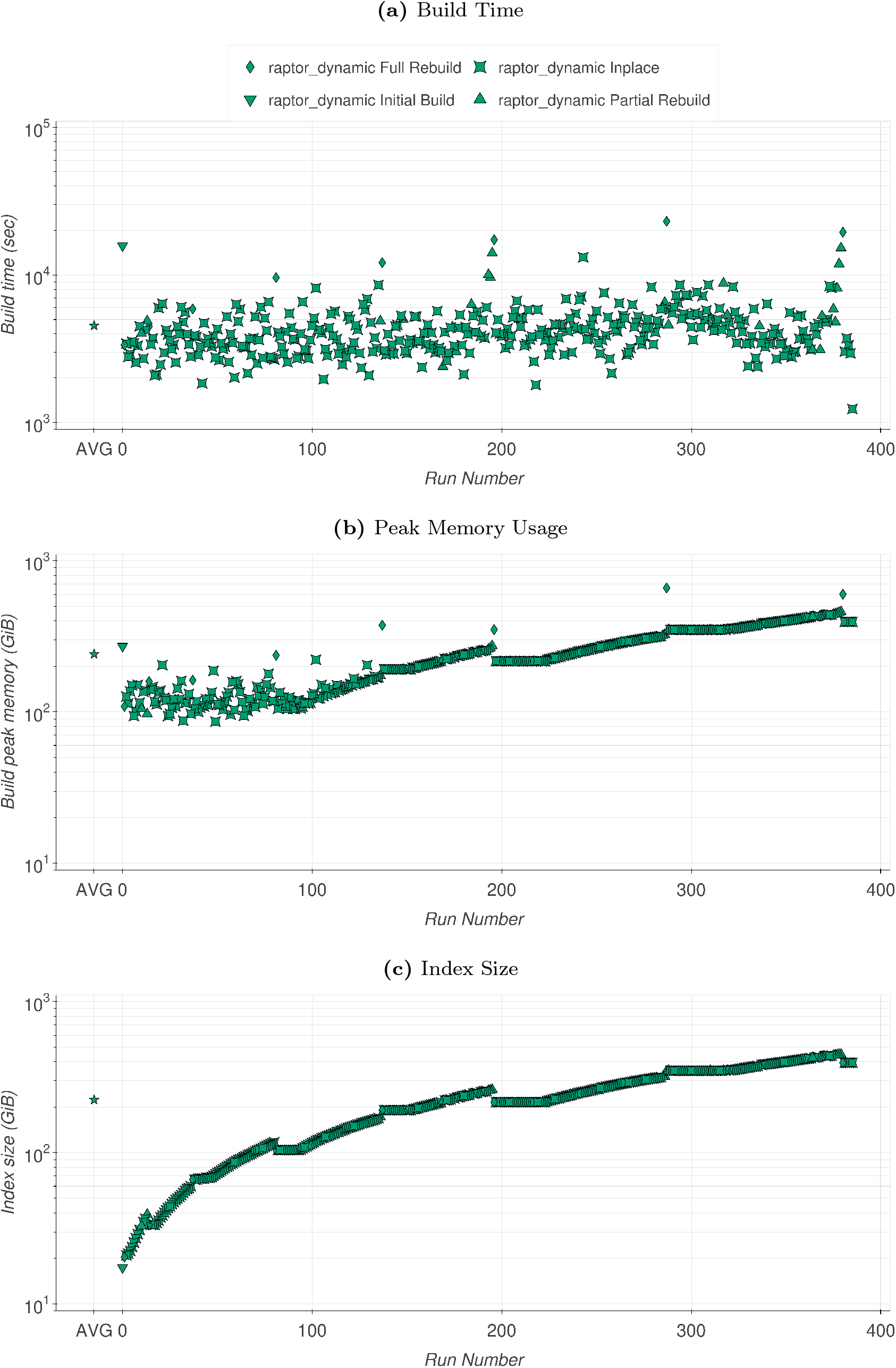
Human RNA 39,400 full samples. Build time in seconds (a), peak memory usage in GiB (b), and index size in GiB (c). The initial build (x=0) contains 1,000 sequences, and each incremental run (x*≥*1) adds 100 sequences. Low-frequency k-mers were removed based on file size.

To assess generalizability, we indexed real microbial genomes (Figure 12) with the parameters of the human RNA data: starting with 10,428 complete archaeal and bacterial RefSeq genomes (from 2024-10-01 to 2025-09-30), adding 13,403 genomes (from 2025-10-01 to 2026-09-30), divided in 53 batches, one for each week of the period. Raptor-Dynamic completed all updates in 1.85 hours, which is 3.9, 10.8, 49, and 109 times faster than Raptor-Static, COBS, Fulgor, and Metagraph, respectively. Only one update required a full rebuild. The final index is 7% larger than that of Raptor-Static, and both are the smallest in this evaluation.

**Figure 12.**
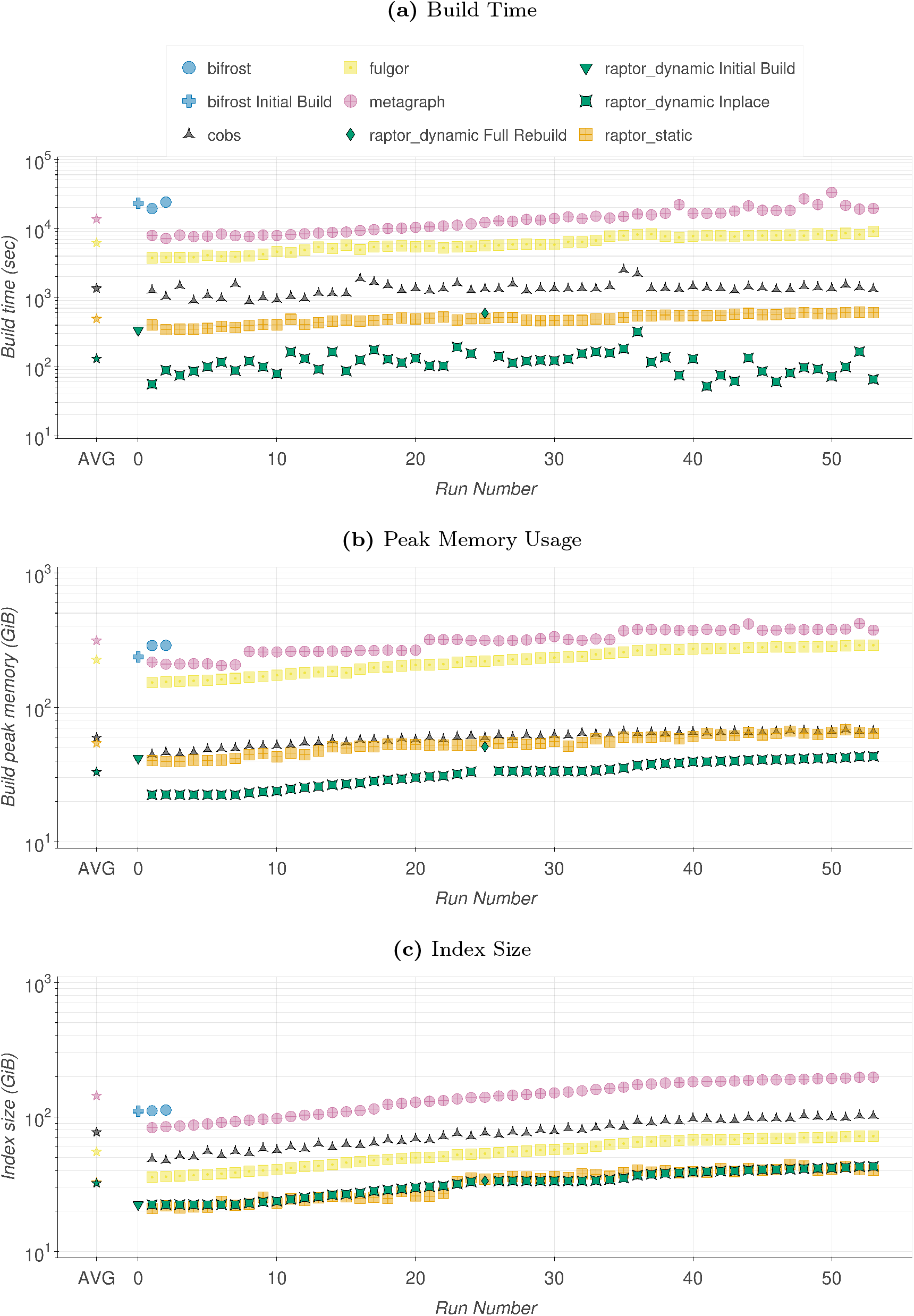
Weekly Batch Update of Real RefSeq Genomes. Build time in seconds (a), peak memory usage in GiB (b), and index size in GiB (c). The initial build (Run Number x=0; Raptor-Dynamic and Bifrost only) contains 10,428 genomes, and each incremental run (x*≥*1) adds the genomes of one week. Bifrost failed after the second update.

**Figure 13.**
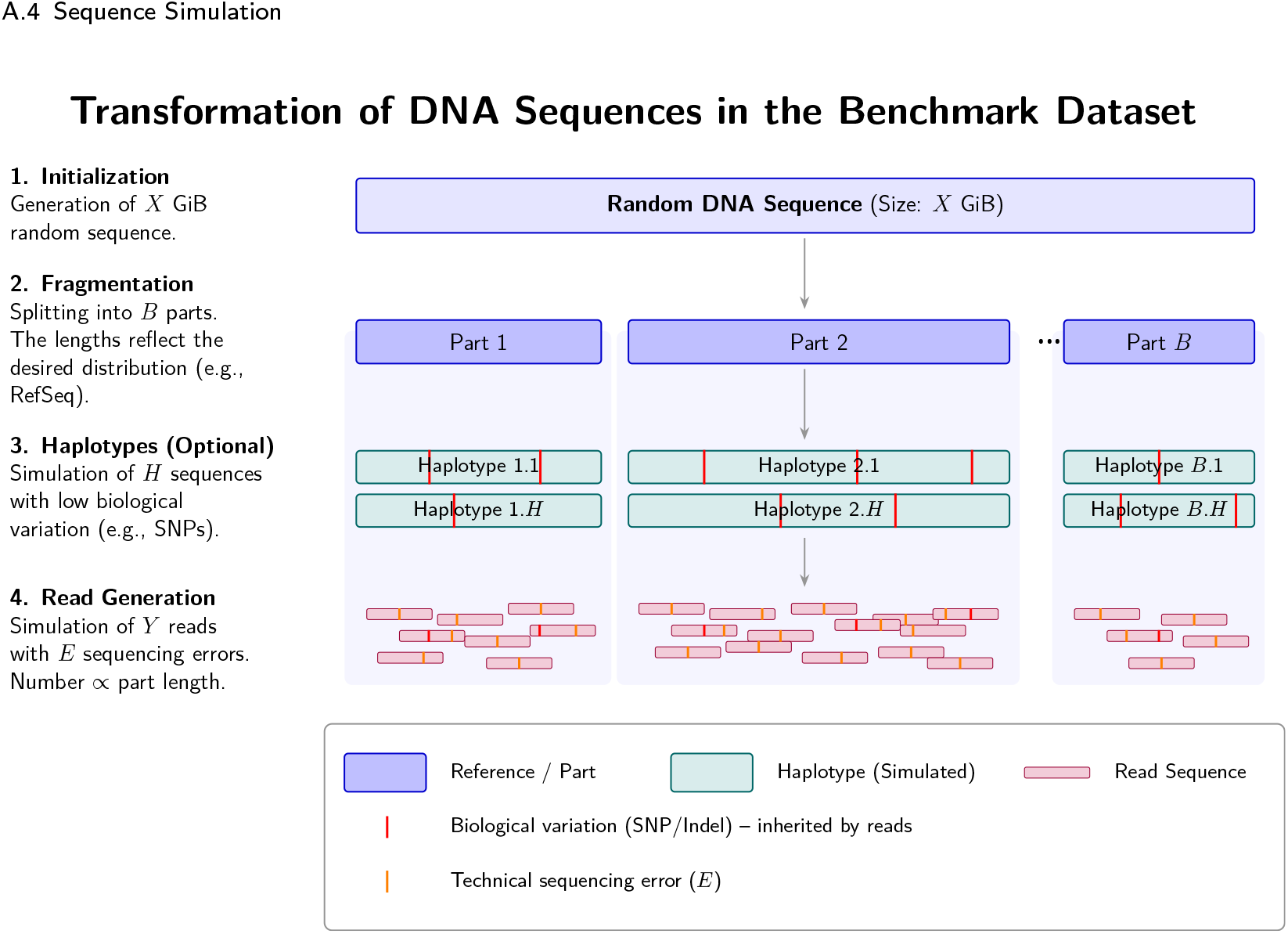
Sequence Simulation. From an initially generated random DNA sequence (*X* GiB), *B* parts are extracted via fragmentation. Optionally, *H* haplotypes are simulated from these reference parts, containing controlled biological variations (e.g., SNPs/Indels, marked in red). Finally, read sequences are simulated from the haplotypes, with the number of reads scaling proportionally to the length of the respective part. The reads inherit the biological mutations and are additionally injected with simulated technical sequencing errors (*E*, marked in orange).

**Figure 14.**
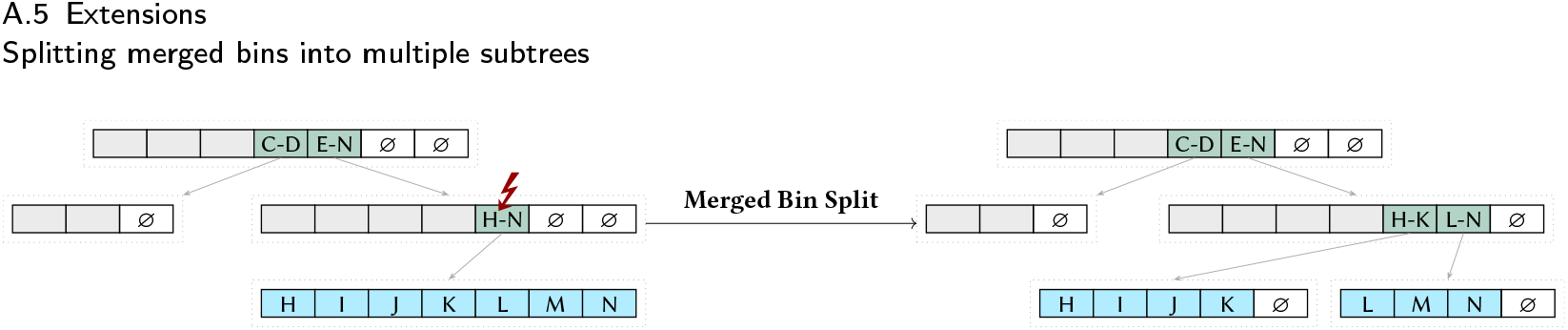
Example merged bin split. Empty bins are denoted by ∅. A gray background is used for present bins that are not relevant for the respective example. A merged bin exceeding the false positive threshold after sample insertion is marked with a f. Instead of a partial rebuild, the affected merged bin is split into two merged bins.

Based on these results, Raptor-Dynamic effectively balances the overhead of incremental updates with high query efficiency. Our findings highlight the potential of dynamic HIBFs for managing rapidly growing sequence datasets, especially when scaling to large-scale genomic indexing and querying.

## 4 Discussion & Conclusion

In this work, we introduced the Dynamic Hierarchical Interleaved Bloom Filter (DHIBF) expanding the functionalities of its predecessor. The state-of-the-art sequencing index previously demonstrated improved scalability, size, and build and query times over competing tools [30]. To be practical for rapidly expanding sequencing archives, we extended it to support dynamic updates, like sample insertions, modifications, and deletions.

Simulated data established the effectiveness of these operations, evaluating scenarios of inserting new sequence content and a mixed workload of insertions and deletions. The dynamic index showed a many-fold improvement in update time over the naive approach of repeatedly rebuilding the index, and its combined update and query time was several times lower than that of the dynamic methods kmindex and Bifrost. Real-world data further underlined the effectiveness of our method. When subsampling 1% of the reads in a human RNA dataset from SRA, the sequential insertion of 5,000 samples was completed in 5.3 hours. This is 23 to 42 times faster than competing tools and still more than twice as fast as our non-dynamic HIBF. Weekly updates of real RefSeq genomes confirmed these results.

This work contributes to the development of more scalable tools for common bioinformatics tasks. The dynamic HIBF might be applied to index large sequence repositories like ENA and SRA, which have grown to a petabases scale and will continue to grow. Our work enables researchers to include a larger portion of the ENA and SRA into existing indices.

While the dynamic HIBF substantially accelerates index updates, its design involves several theoretical trade-offs and structural constraints: Reserving empty technical bins and opportunistically resizing IBFs introduces a space overhead compared to the static HIBF; however, with default settings, this overhead is small on average (2% to 4%), although a full rebuild can temporarily yield a considerably larger index (Figure 6c, Figure 7c, Figure 10c).

- Relaxing the false positive rate threshold in merged bins via 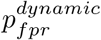 theoretically increases the probability of false-positive traversals into lower-level IBFs, though we observed no noticeable impact on empirical query performance (Table 2).
- Because standard Bloom filters do not support single-element deletion, deleting a sample clears its technical bins, whereas its *k*-mers remain in parent merged bins unless all child bins of a merged bin are deleted or a rebuild is triggered.
- Split samples currently require contiguous empty technical bins, meaning that scattered empty bins resulting from prior deletions cannot be combined to store a split sample without resizing or rebuilding.
- Because our approach relies on amortized partial and global rebuilding, individual update times and memory usage are not worst-case bounded: while most updates execute in-place using minimal RAM, occasional full rebuilds temporarily require the higher runtime and peak memory footprint of static index construction.

In conclusion, we believe that the DHIBF constitutes an important step forward for working with truly large data sets that change slowly over time.

Lastly, we want to mention that there are various ideas for extensions that could further improve the performance of the dynamic HIBF which we sketch in the appendix (Appendix A.5).

## Supporting information

Tracked Changes

## Declarations

## Abbreviations

BF: Bloom Filter
IBF: Interleaved Bloom Filter
HIBF: Hierarchical Interleaved Bloom Filters
DHIBF: Dynamic Hierarchical Interleaved Bloom Filters
FPR: False Positive Rate
UB: User Bin
TB: Technical Bin
EB: Empty Bin

## Ethics approval and consent to participate

Not applicable

## Consent for publication

Not applicable

## Availability of data and materials

The list of accessions used for the RNA-Seq data set is available at https://github.com/splatlab/mantis/blob/mergeMSTs/experiments/humanRNA_40k.accessions. The simulated data sets as well as raw results are available at https://zenodo.org/records/21689968. Interactive benchmark results are available at https://docs.seqan.de/dynamic/, the benchmark workflow at https://github.com/seqan/raptor/tree/main/util/dynamic.

## Authors’ contributions

KR initiated the project and the main idea for the DHIBF. MW devised in her MSc thesis a first prototype for a DHIBF. ES integrated the viable methods into the main HIBF branch. VP conducted the experiments and wrote the Results section. All authors read and approved the final manuscript.

## Acknowledgements

Not applicable

## Availability and Requirements

Project name: Raptor

Project home page: https://github.com/seqan/raptor

Operating system(s): Linux, macOS, BSD

Programming language: C++

Other requirements: For compiling: GCC *≥* 14, or Clang *≥* 19.

No additional runtime requirements.

License: BSD-3-Clause

Any restrictions to use by non-academics: No restriction

## Appendix A: Appendix

### A.1 Hardware specifications

#### CPU

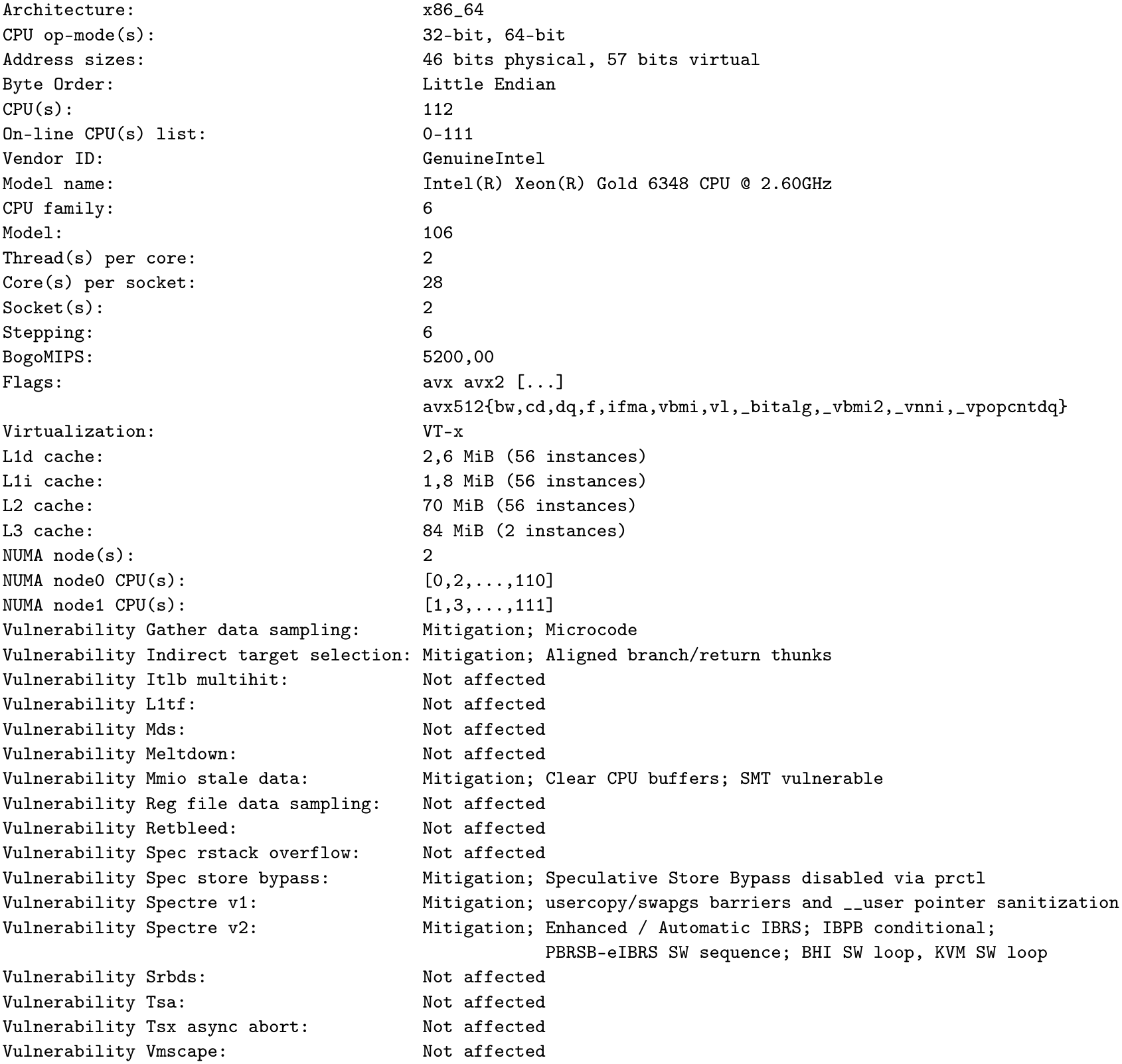

#### Kernel

`Linux 6.1.0-43-amd64 #1 SMP PREEMPT_DYNAMIC Debian 6.1.162-1 (2026-02-08) x86_64 GNU/Linux`

### A.2 Supplemental Data

The source code is written in C++23 using the SeqAn Library [36] and available at https://github.com/seqan/raptor. Additional resources for this project, including sample metadata and analysis scripts, are available in the latter GitHub repository.

### A.3 Existing tools

### A.4 Sequence Simulation

### A.5 Extensions

#### Splitting merged bins into multiple subtrees

During the rebuilding process as described in Section 2.5, the complete subtree needs to be rebuilt. A feasible alternative could be to split the affected merged bin into two or more merged bins (Appendix Figure 14). This reduces the number of bins that need to be accounted for during the rebuild. Additionally, it would allow using bit shift or bin-deletion operations to reuse existing lower-level IBFs. For example, in Appendix Figure 14, the IBF containing H-K after splitting could also be created by removing L-N from the original IBF containing H-N. Consequently, the H-K IBF would then have 3 empty bins instead of 1.

#### Bin adjacency

In the current implementation, split samples are required to be placed in adjacent bins. Empty bins that are not adjacent can only be used for single bins. To increase flexibility when placing new samples, the algorithm could be modified to allow single samples to be placed in non-adjacent bins within an IBF. However, this would also require additional bookkeeping. Alternatively, functionality could be added to restore empty bin adjacency by shifting bits.

#### Batch insertions

As of now, multiple samples can be provided as input at once. However, internally, updates occur sequentially, which can slow down batch insertion. The process may be sped up by prioritizing the largest samples first, to avoid unnecessary small rebuild operations that are triggered by smaller samples. A more informed approach would involve creating a new layout for the batch and then potentially inserting merged bins.

#### Accounting for update and query frequencies

Another extension could lead to further advancements in overall usability. By tracking the frequency of updates and queries to samples stored in the DHIBF, we could achieve more efficient updates and faster queries, particularly for frequently searched samples like *E. coli*.. We can draw inspiration from data structures like splay trees, which update the tree after querying by placing the queried entry higher in the tree. For the DHIBF, this would translate to reserving more empty bins for parts of the index that are often updated and placing heavily queried parts higher in the hierarchy when rebuilding.

## Footnotes

[1] 43,090 complete genomes of archaea and bacteria, as of 2024-08-26

[2] https://github.com/seqan/raptor/tree/main/util/simulation

[3] https://github.com/splatlab/mantis/blob/mergeMSTs/experiments/humanRNA_40k.accessions

[4] https://github.com/ratschlab/metagraph/issues/574

[5] https://github.com/Malfoy/PAC/issues/6

[6] https://github.com/splatlab/mantis/issues/24

[7] https://github.com/tlemane/kmindex/issues/19

[8] https://github.com/pmelsted/bifrost/issues/98

