## Supplementary material for "Dynamic Hierarchical Interleaved Bloom Filter: An Updatable Index for Large-Scale Fast Sequence Search": Tracked Changes

While many approaches for dynamic updates exist (see Section 1.3), ~~we often found the practical implementation to be dissatisfying for six reasons~~existing implementations exhibit six primary limitations in practice:

**VariMerge** [27] extends the Vari data structure using a divide-and-conquer approach, splitting large datasets into smaller sets and merging their cdBGs. In addition, VariMerge supports sample insertions.

**Table 1 Algorithmic comparison of multi-set sequence indexing tools.** Overview of representative colored de Bruijn graph and sequence aggregation indices, detailing their support for dynamic insertion and deletion operations along with their core indexing mechanisms. An expanded version of this table can be found in ~~Appendix 3~~Appendix Table 3.

| Tool | Insert | Delete | Description | Reference |
|---|---|---|---|---|
| Bifrost | ✓ | ∼ | Builds compacted colored de Bruijn graphs using a minimizer-based hash table mapped to unitigs. Deletion only via API. | [16] |
| BIGSI | ✓ | × | Stacks identical Bloom filters for each dataset into a column-major bitmatrix queried via a hash table. Index update requires modifying every key in the index. | [28] |
| COBS | × | × | A compact bit-sliced signature index linking an inverted index with Bloom filters, grouping files of similar sizes to save space. | [17] |
| Dynamic Mantis | ✓ | × | Replaces RRR compression with an MST-based compression of color classes. Uses the Bentley-Saxe transformation to merge indexes iteratively. | [29] |
| Fulgor | × | × | Maps ccdBG unitigs to colors using an order-preserving SSHash dictionary, maintaining unitigs in exact color order to compress integer arrays. | [18] |
| HowDe-SBT | ✓ | ✓ | Optimizes SBTs by stripping out redundant "non-informative" bits, drastically shrinking the tree's memory footprint. | [19] |
| kmindex | ✓ | × | Builds partitioned, inverted one-hash Bloom filter matrices aligned by minimizers. | [20] |
| MetaGraph Default | × | × | Uses a BOSS-encoded SuccinctDBG and highly compressed annotation matrices (e.g., Multi-BRWT). | [24] |
| MetaGraph Dynamic | ✓ | ✓ | Adapts the static MetaGraph approach by backing the core graph vectors with dynamic, cache-efficient B-trees. | [24] |
| PAC | ✓ | × | Partitions k-mers by minimizers, structuring Aggregated Bloom filters into Bloom Comb Trees. | [25] |
| SeqOthello | ✓ | × | Relies on Othello minimal perfect hashing to map k-mers to their corresponding occurrence profiles. Updates require a merge and hash reconstruction. | [26] |
| VariMerge | ✓ | × | Extends the Vari architecture to support the efficient merging of distinct, updatable cdBG representations. | [27] |

figure changed

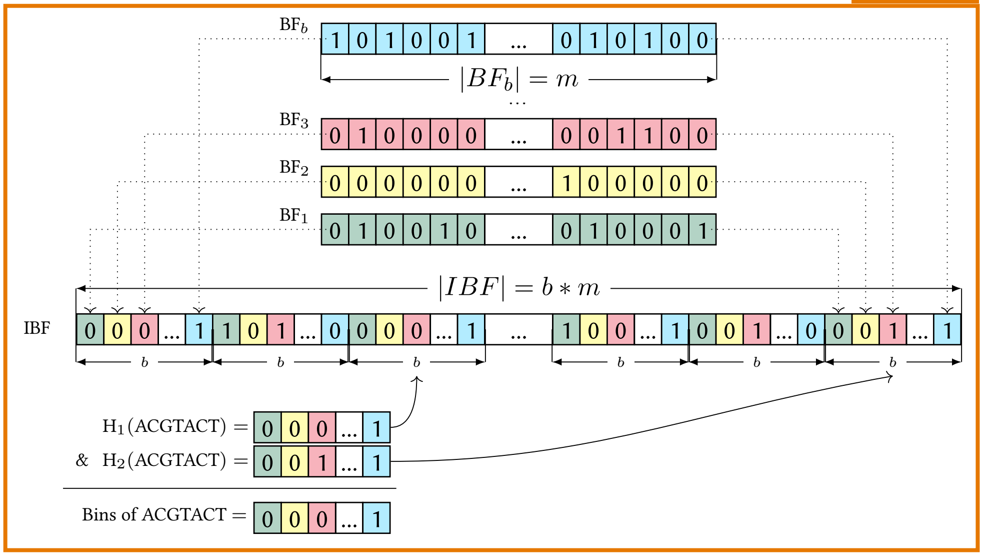


**Figure 1 Structuring and querying an Interleaved Bloom Filter (IBF).** Differently colored BFs of bin size ~~$n$~~ $m$ for $b$ samples (shown on the top) are interleaved and yield an IBF of size ~~$b \times n$~~ $b \times m$. In the example, $h = 3$ different hash functions are used to store and query a $k$-mer. When inserting a $k$-mer from bin $B_i$ into the IBF, all $h$ hash functions are computed, pointing to the positions of the corresponding $h$ sub-bitvectors, in which the $i^{\text{th}}$ bits are set to 1. When querying a $k$-mer, $h$ sub-bitvectors are retrieved and combined using a bitwise AND to obtain the binning bitvector of length $b$, where a 1 indicates the presence of the $k$-mer in the corresponding bin. The figure has been adapted from [32].

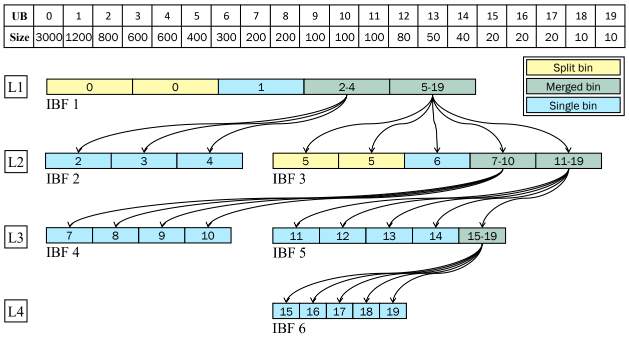


**Figure 2 The hierarchical index structure of the HIBF** This exemplary HIBF of 19 numbered user bins (UB) consists of four levels. The top-level (L1) has always a single IBF (IBF-1) with exactly $t_{max}$ technical bins, in this case 5. This IBF stores the complete dataset using notably less space than a regular IBF because of its split and merged bins (yellow and green, respectively). Each IBF displays boxes representing its technical bins, which store the $k$-mer content of the labelled user bins. For instance, in IBF-1, UB-1 is split and stored in two technical bins, UB-2 in a single technical bin, and UB-2 to 4 are merged into one technical bin. Note that the schematic does not depict that in reality these technical bins are interleaved. IBFs in lower levels each correspond to a merged bin in a previous level. For instance, IBF-2 and IBF-3 on the second level (L2) correspond to the first and second merged bin of IBF-1. Note that the IBFs in the layout form a tree, where the root is the top-level IBF and the leaves consist of IBFs without merged bins. The figure was adapted from [30].

When a new sample is to be inserted in the index, its $k$-mers are extracted and counted. Then, we compute the capacity $n$ of each IBF, i.e., the maximum number

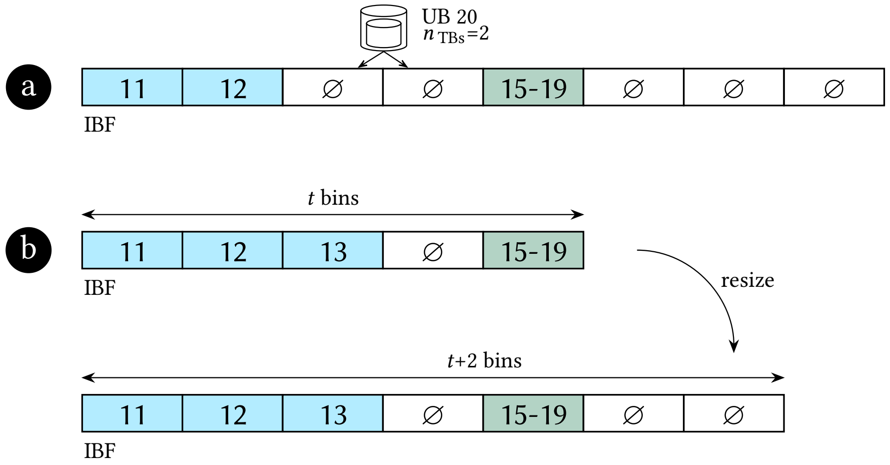


**Figure 3 Selecting IBFs for sample insertions**. Empty bins are denoted by ∅. After selecting an IBF, the algorithm scans an auxiliary table from left to right to find adjacent empty bins within the IBF. In example **(a)**, there are enough ($\geq n_{\text{TB}}$) adjacent empty bins to accommodate the new sample. In case insufficient adjacent empty bins are available (**b**), a resize operation is executed. Enough empty bins (EB) are added such that the new sample can be accommodated. However, the number of technical bins is not allowed to exceed $t_{max} + 64$.

of elements each IBF can accommodate while still maintaining the FPR constraints (Equation 2).

$$\begin{aligned}
& p_{fpr} = (1 - e^{-hn/m})^h && (1)\\
\Leftrightarrow\quad & \sqrt[h]{p_{fpr}} = 1 - e^{-hn/m}\\
\Leftrightarrow\quad & e^{-hn/m} = 1 - \sqrt[h]{p_{fpr}}\\
\Leftrightarrow\quad & -\frac{hn}{m} = \ln\left(1 - \sqrt[h]{p_{fpr}}\right)\\
\Leftrightarrow\quad & -hn = m \ln\left(1 - \sqrt[h]{p_{fpr}}\right)\\
\Leftrightarrow\quad & n = -\frac{m \ln\left(1 - \sqrt[h]{p_{fpr}}\right)}{h}
\end{aligned}$$

$$\begin{aligned}
& p_{fpr} \approx (1 - e^{-hn/m})^h && (1)\\
\Leftrightarrow\quad & \sqrt[h]{p_{fpr}} \approx 1 - e^{-hn/m}\\
\Leftrightarrow\quad & e^{-hn/m} \approx 1 - \sqrt[h]{p_{fpr}}\\
\Leftrightarrow\quad & -\frac{hn}{m} \approx \ln\left(1 - \sqrt[h]{p_{fpr}}\right)\\
\Leftrightarrow\quad & -hn \approx m \ln\left(1 - \sqrt[h]{p_{fpr}}\right)\\
\Leftrightarrow\quad & n \approx -\frac{m \ln\left(1 - \sqrt[h]{p_{fpr}}\right)}{h}
\end{aligned} \tag{2}$$

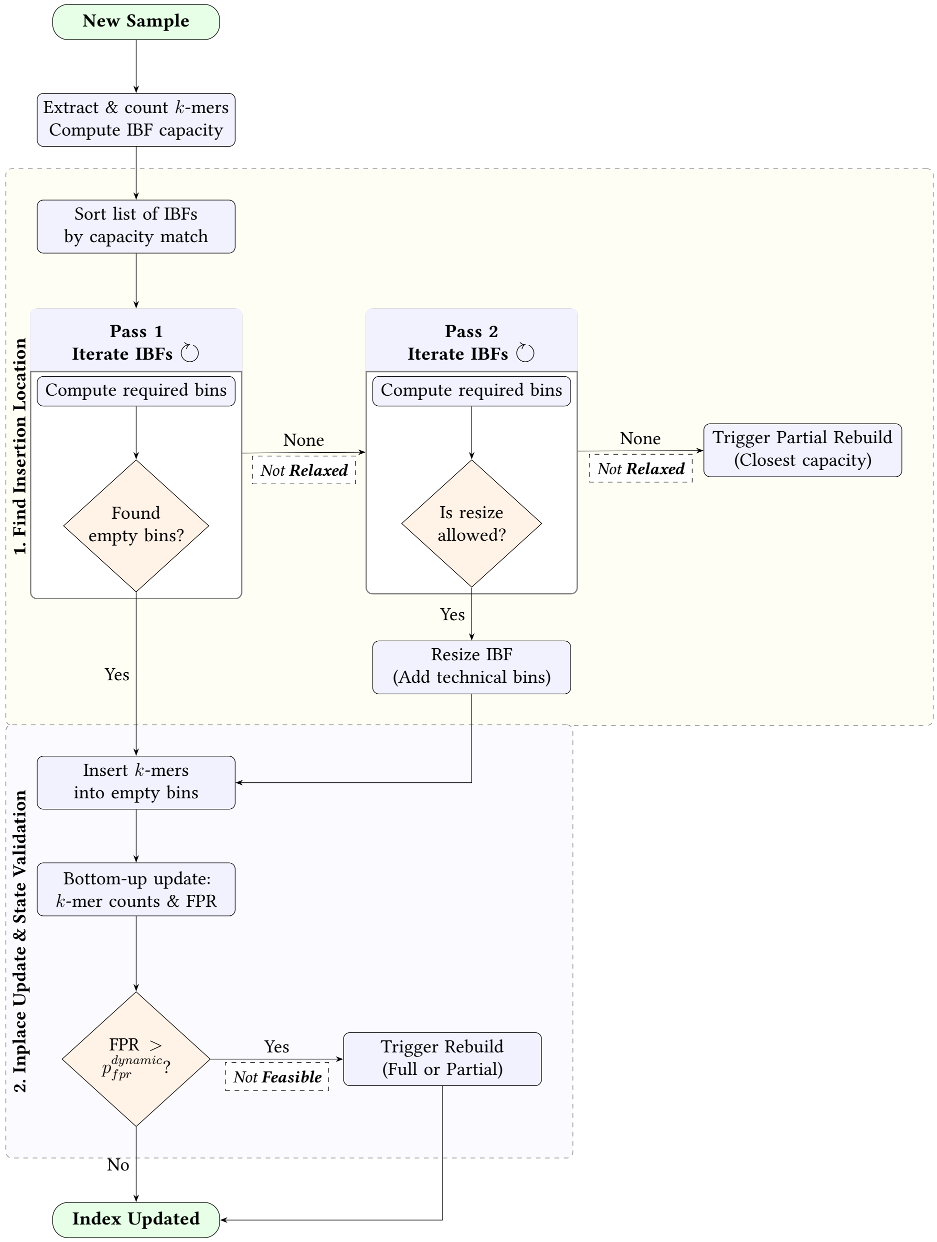


**Figure 4 Schematic overview of the dynamic HIBF sample insertion**. The process ensures the index remains in a *relaxed* and *feasible* state by searching for adjacent empty bins, opportunistically resizing IBFs to prevent immediate bottlenecks, and evaluating dynamic FPR thresholds ($p_{fpr}^{dynamic}$) to trigger necessary rebuilds.

#### *2.4.1 Finding the Insertion Location*

Next, we search for an IBF to insert the new sample into. We sort the list of IBFs by how closely their capacity matches the $k$-mer count and evaluate each IBF until we find a suitable location. The evaluation begins by computing the number of technical bins needed to store the new sample. If the current IBF has sufficient consecutive empty bins available, it is selected as the insertion location (Figure 5a). In case no IBF is suitable, we repeat the evaluation process with the added possibility to resize an IBF (Figure 5b). Resizing an IBF allows us to add more technical bins to an existing IBF without constructing it from scratch. This is much faster than (partially) rebuilding the full index, albeit only when done sparsely. We allow resizing up to $t_{max} + 64$ technical bins. Resizing is not allowed in three cases: 1) the IBF has already been resized, 2) the IBF is the root IBF, 3) the IBF has been part of a partial rebuild. If this second search is unsuccessful, a partial rebuild for the IBF with the closest capacity is triggered. The partial rebuild will make the IBF and its children ineligible for future resizing.

#### 2.4.2 Inplace Update

The $k$-mers are inserted into the empty bins while updating associated auxiliary metadata. If the target IBF was not the root IBF, this process is repeated for the parent IBF until we arrive at the root IBF.

While inserting $k$-mers, we also keep track of the number of elements that have been inserted into a technical bin. Note that the number of elements a technical bin (or, conceptually, a Bloom filter) contains is most likely different from the number of $k$-mers inserted due to potential hash collisions. Put differently, adding a $k$-mer to a Bloom filter might not set any additional bits. Hence, the number of elements in the Bloom Filter is always less than or equal to the number of $k$-mers inserted. Using the exact number of elements contained in a Bloom filter allows for an exact computation of the false positive rate for any technical, and especially merged, bins. If the allowed FPR is exceeded for any IBF while inserting, we remember the IBF's location to later trigger a partial rebuild and continue with the insertion.

While inserting $k$-mers, we also track the effective number of elements added to each technical bin to monitor its false positive rate (FPR). Specifically, we only increment the bin's element counter if inserting the $k$-mer sets at least one previously unset bit, i.e., the incoming $k$-mer is not reported as present by the Bloom filter. Because a Bloom filter cannot distinguish between a duplicate $k$-mer and a false-positive hash collision of a novel $k$-mer, this counter represents the number of bit-modifying insertions rather than the exact number of distinct $k$-mers. However, because $k$-mers that already test positive do not set any additional bits, they do not increase the bitvector's saturation or degrade the actual FPR. Substituting this effective element count $n$ into Equation 1 therefore provides a close estimate of the technical bin's current FPR, particularly for merged bins where samples share $k$-mers, without overcounting redundant insertions. If the estimated FPR exceeds the allowed threshold for any IBF during insertion, we record the IBF's location to trigger a partial rebuild once the insertion completes. The FPR threshold depends on the type of technical bin the insertion occurs in and, for merged bins, on its level (depth within the tree). Split bins have to conform to $p_{fpr}^{max}$. For merged bins, we use the merged-bin-specific FPR $p_{fpr}^{merged}$ with a level-dependent relaxation factor:

$$p_{fpr}^{dynamic}(level) = \left(1 + \frac{1}{(level + 1)^2}\right) \cdot p_{fpr}^{merged} \tag{3}$$

The *level* is 0 for the root IBF and increases by one for each merged bin encountered along the path to an IBF. The dynamic FPR $p_{fpr}^{dynamic}$ accounts for the presence of more children on higher levels while not precluding partial rebuilds of lower-level subtrees. Note that some form of relaxation factor is necessary for dynamization. When an HIBF is constructed, it is likely that the fullest bin is a merged bin, in which case it $p_{fpr}^{merged}$ is used to determine the required bin size. As this is a strict lower bound for the bin size, any insertion into the merged bin would immediately trigger a rebuild.

After selecting an IBF, the algorithm looks for empty bins within this IBF using an auxiliary table (Figure 3, Figure 5a). If there are not enough adjacent empty bins available, a resize and possibly a rebuild operation (Figure 5c) will be triggered.

Finally, the $k$-mers are hashed into the empty bins, and auxiliary tables with FPRs and $k$-mer counts are updated for the bins along the insertion path. The insertion into the hierarchical index is done bottom-up and the auxiliary tables are updated simultaneously. To update the $k$-mer counts of the merged bins, we check for each new $k$-mer if it was already present when inserting it. Only the number of $k$-mers that were not already present is added to the count. If any of the FPRs exceed the FPR threshold, a rebuild operation is triggered (Figure 5c).

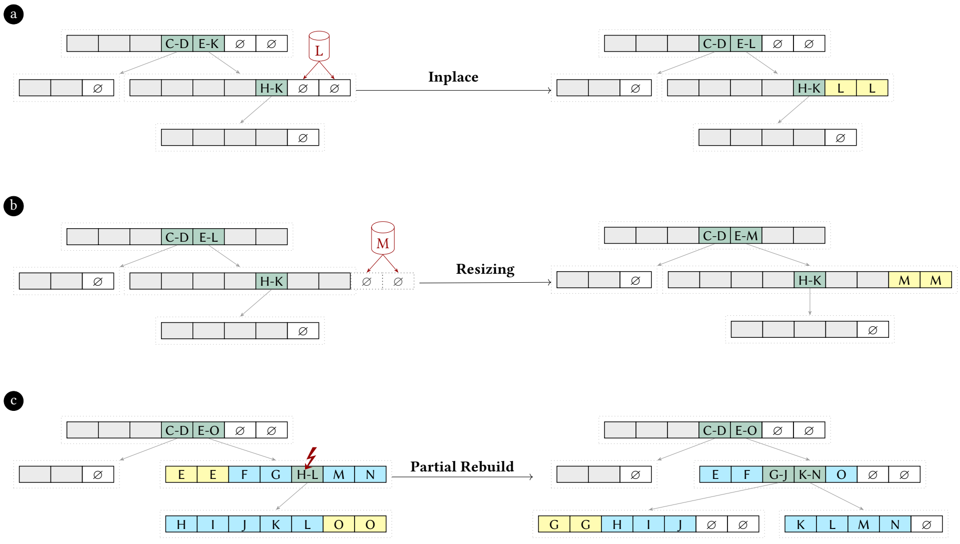


**Figure 5 Sample insertion examples.** Empty bins are denoted by ∅. A gray background is used for present bins that are not relevant for the respective example. A merged bin exceeding the false positive threshold after sample insertion is marked with a ↯. The left-hand side shows the initial index status, while the right-hand side shows the index after insertion. a) depicts an inplace insertion of user bin $L$. b) shows how the user bin $M$ is inserted by resizing an IBF. c) illustrates a partial rebuild triggered by exceeding $p_{fpr}^{dynamic}$ at the merged bin $H - L$.

[1]43,090 complete genomes of archaea and bacteria, as of 2024-08-26
[2]https://github.com/seqan/raptor/tree/main/util/simulation
[3]https://github.com/splatlab/mantis/blob/mergeMSTs/experiments/humanRNA_40k.accessions
[4]https://github.com/ratschlab/metagraph/issues/574
[5]https://github.com/Malfoy/PAC/issues/6
[6]https://github.com/splatlab/mantis/issues/24
[7]https://github.com/tlemane/kmindex/issues/19

it on the simulated data only, where Raptor does the same (it uses minimizers with a window size of 24 on real data). On real data, Bifrost crashed after at most two updates[8]. We also considered kmindex [20], which supports index extension, but did not include it in this evaluation since its utility is limited by a maximum k-mer size of 15,and it requires manual pre-computation of index sizes (a non-trivial task already automated in other tools).

When evaluating the results of querying the simulated reads ~~with two errors~~ for both even and RefSeq datasets, ~~all tools~~ Raptor-Dynamic, Raptor-Static, COBS, Metagraph, and Bifrost achieve perfect accuracy ~~besides Fulgor, which~~ in all runs, with identical results for both Raptor variants. kmindex only has up to 9 false positives per run on the RefSeq-like dataset (0.003% overall). Fulgor consistently had a significant number of false positives and false negatives, around 4.8% and 1.8% on average for all batches, respectively. The number of false positives gradually declines until it reaches 0 after the last batch, when all samples are in the index.

**Table 2** **Build and query times on the simulated datasets.** Times in seconds over runs 1–90: build time per update and time to query all reads at the given number of indexed samples (runs 10, 30, 50, 70, and 90). Tools updating an index are listed first.

| | Build | | Query, by number of indexed samples | | | | | Query |
|---|---|---|---|---|---|---|---|---|
| Tool | mean | median | 20,000 | 40,000 | 60,000 | 80,000 | 100,000 | mean |
| *Evenly simulated* | | | | | | | | |
| Raptor-Dynamic | 17.5 | 2.8 | 0.34 | 0.64 | 0.63 | 0.80 | 1.87 | 0.67 |
| kmindex | 29.2 | 29.5 | 45.9 | 131 | 154 | 242 | 228 | 180 |
| Bifrost | 465.8 | 447.1 | 146 | 290 | 503 | 603 | 999 | 422 |
| Raptor-Static | 247.1 | 194.3 | 0.37 | 0.65 | 0.66 | 0.92 | 1.31 | 0.76 |
| COBS | 10.4 | 11.4 | 40.1 | 81.9 | 112 | 124 | 172 | 107 |
| Fulgor | 222.9 | 216.7 | 1.12 | 2.03 | 1.88 | 2.62 | 2.23 | 2.21 |
| Metagraph | 174.5 | 146.1 | 72.1 | 226 | 477 | 799 | 1122 | 465 |
| *RefSeq-like simulated* | | | | | | | | |
| Raptor-Dynamic | 33.0 | 7.7 | 0.33 | 0.58 | 0.75 | 0.85 | 1.15 | 0.68 |
| kmindex | 20.1 | 20.0 | 32.3 | 64.4 | 97.5 | 132 | 167 | 93.6 |
| Bifrost | 507.0 | 501.9 | 123 | 449 | 449 | 608 | 757 | 437 |
| Raptor-Static | 425.6 | 331.0 | 0.50 | 0.80 | 0.77 | 1.36 | 1.08 | 0.76 |
| COBS | 11.3 | 11.6 | 66.3 | 81.2 | 113 | 117 | 157 | 104 |
| Fulgor | 224.8 | 221.0 | 1.14 | 1.55 | 2.29 | 2.28 | 2.34 | 2.08 |
| Metagraph | 154.1 | 134.0 | 74.8 | 211 | 484 | 755 | 1107 | 477 |

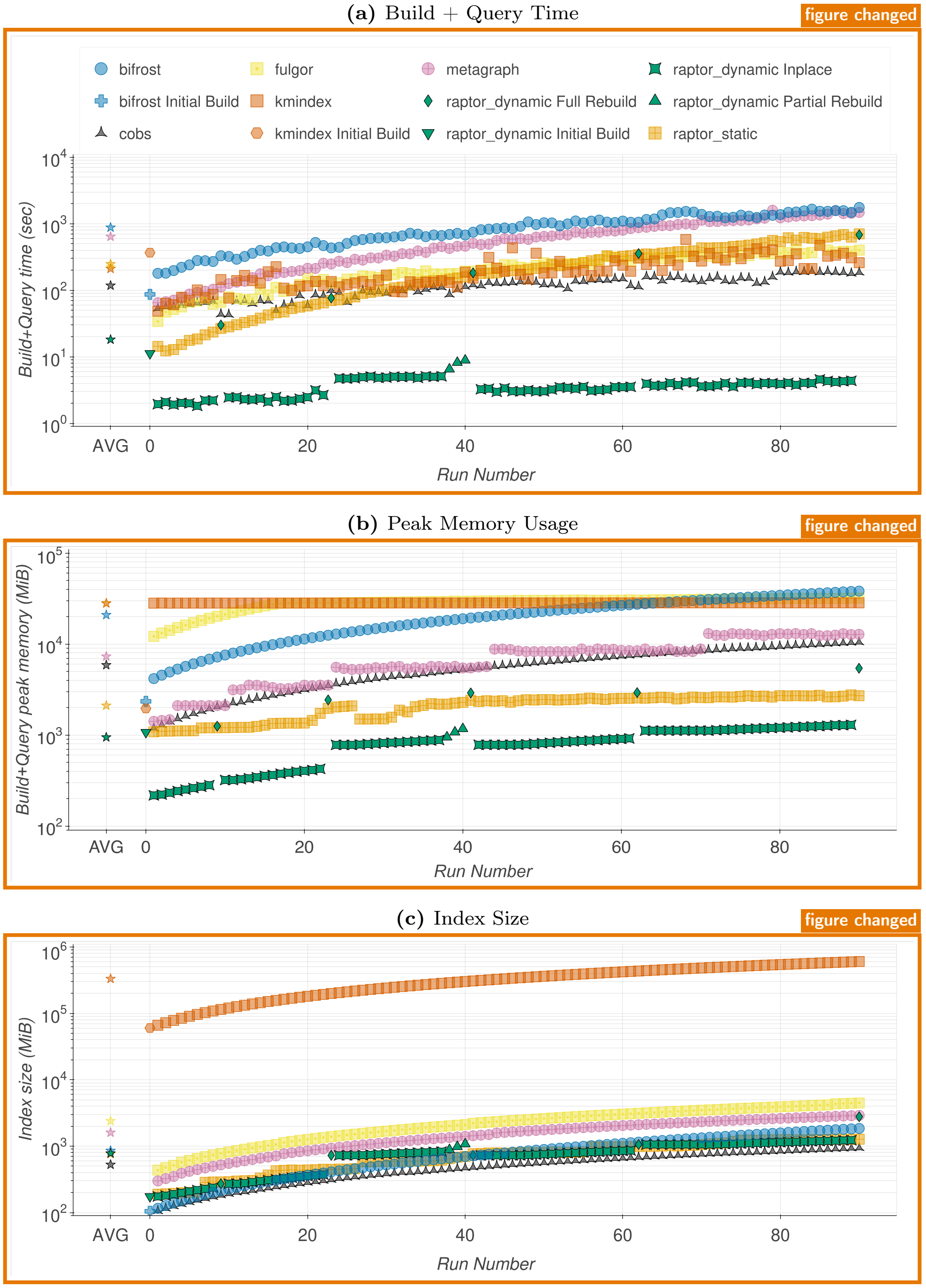


**Figure 6** ~~Simulated Batch Even.~~ Batch Update of Evenly Simulated Dataset. Build + Query time in seconds (a), peak memory usage in MiB (b), index size in MiB (c). The initial build (Run Number x=0; Raptor-Dynamic, ~~Raptor~~ kmindex, and Bifrost only) contains 10,000 sequences, and each incremental run (x≥1) adds 1,000 sequences. Leftmost points are the average among all data points for each tool, color-based.

**(a)** Build + Query Time

figure changed

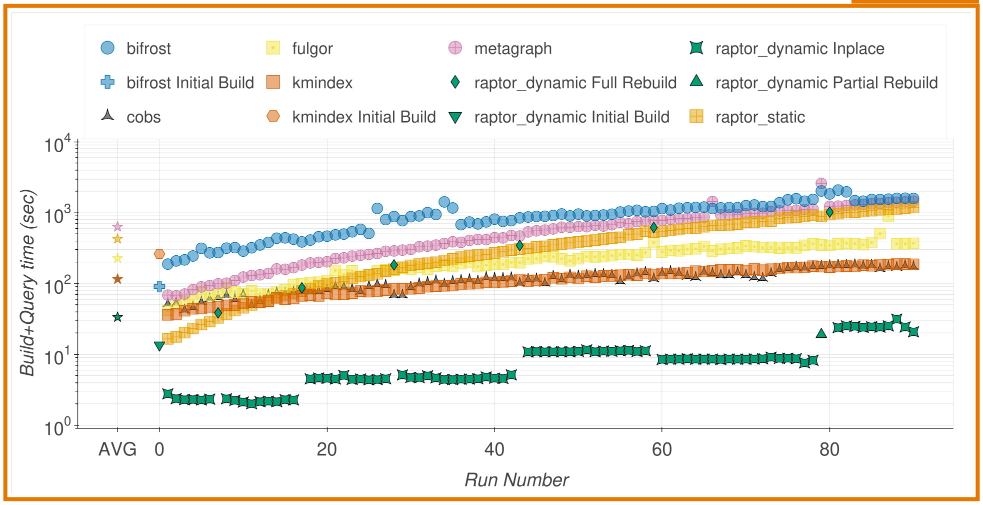


**(b)** Peak Memory Usage

figure changed

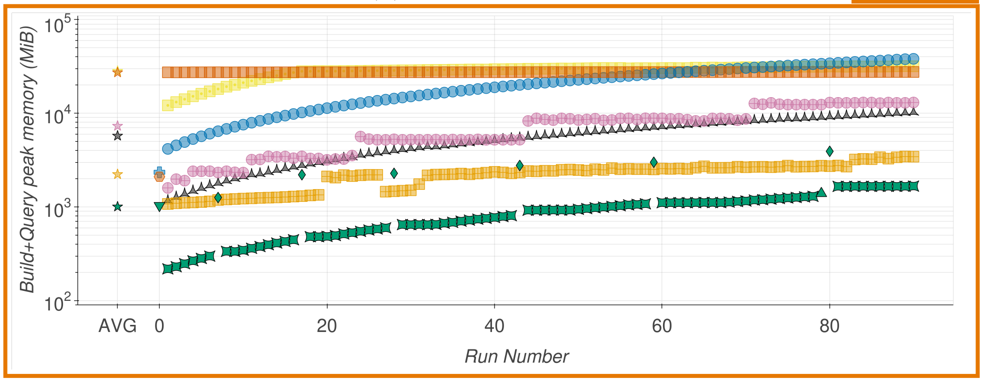


**(c)** Index Size

figure changed

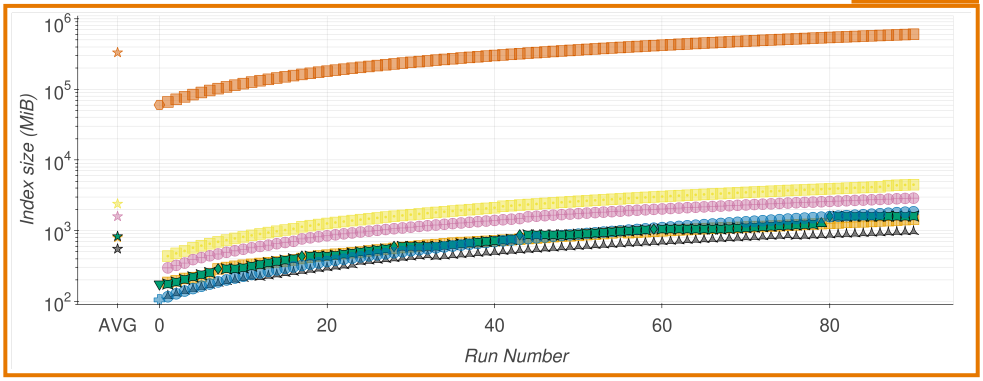


**Figure 7** ~~Simulated Batch RefSeq.~~ Batch Update of RefSeq-like Simulated Dataset. Build + Query time in seconds (a), peak memory usage in MiB (b), index size in MiB (c). The initial build (Run Number x=0; Raptor-Dynamic, ~~Raptor~~ kmindex, and Bifrost only) contains 10,000 sequences, and each incremental run (x≥1) adds 1,000 sequences. Leftmost points are the average among all data points for each tool, color-based.

**(a)** Build + Query Time

figure changed

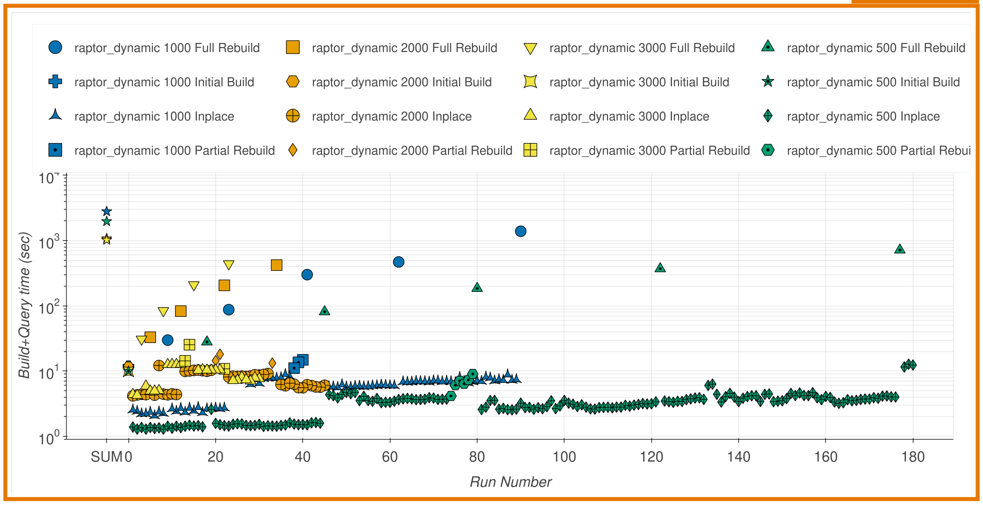


**(b)** Peak Memory Usage

figure changed

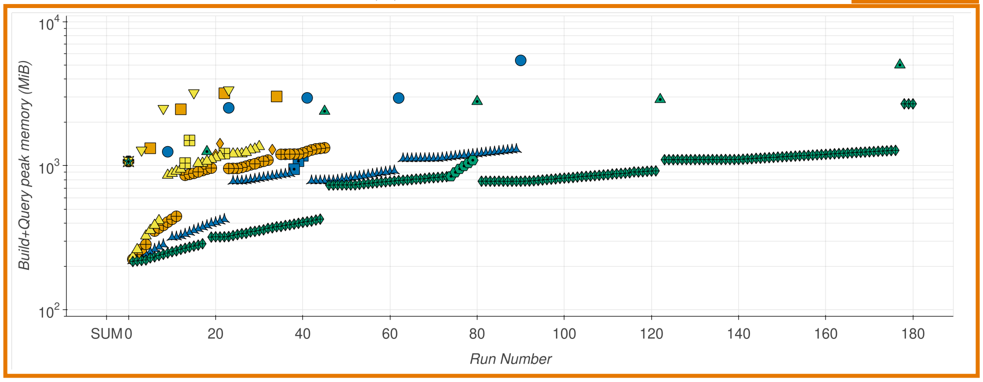


**(c)** Index Size

figure changed

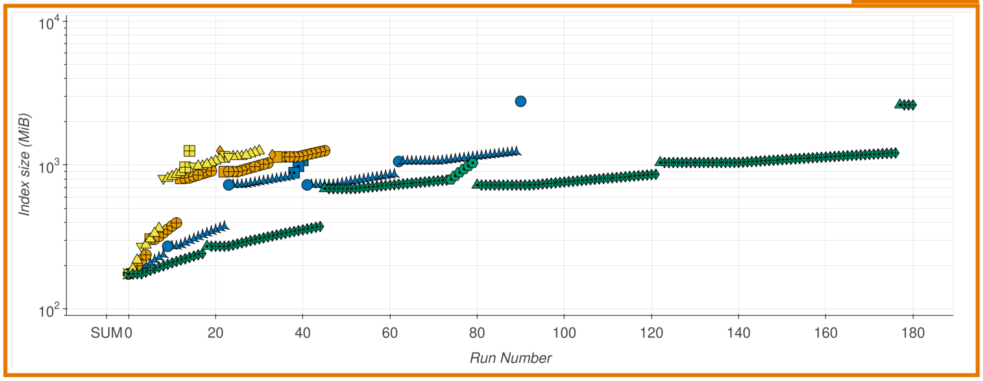


**Figure 8** **Batch Update of Evenly Simulated Dataset with Different Batch Sizes.** Build + Query time in seconds (a), build peak memory usage in MiB (b), index size in MiB (c). The initial build (Run Number x=0) contains 10,000 sequences, and each incremental run (x≥1) adds 500, 1,000, 2,000, or 3,000 sequences. Leftmost points are the sum among all data points for each variation, color-based.

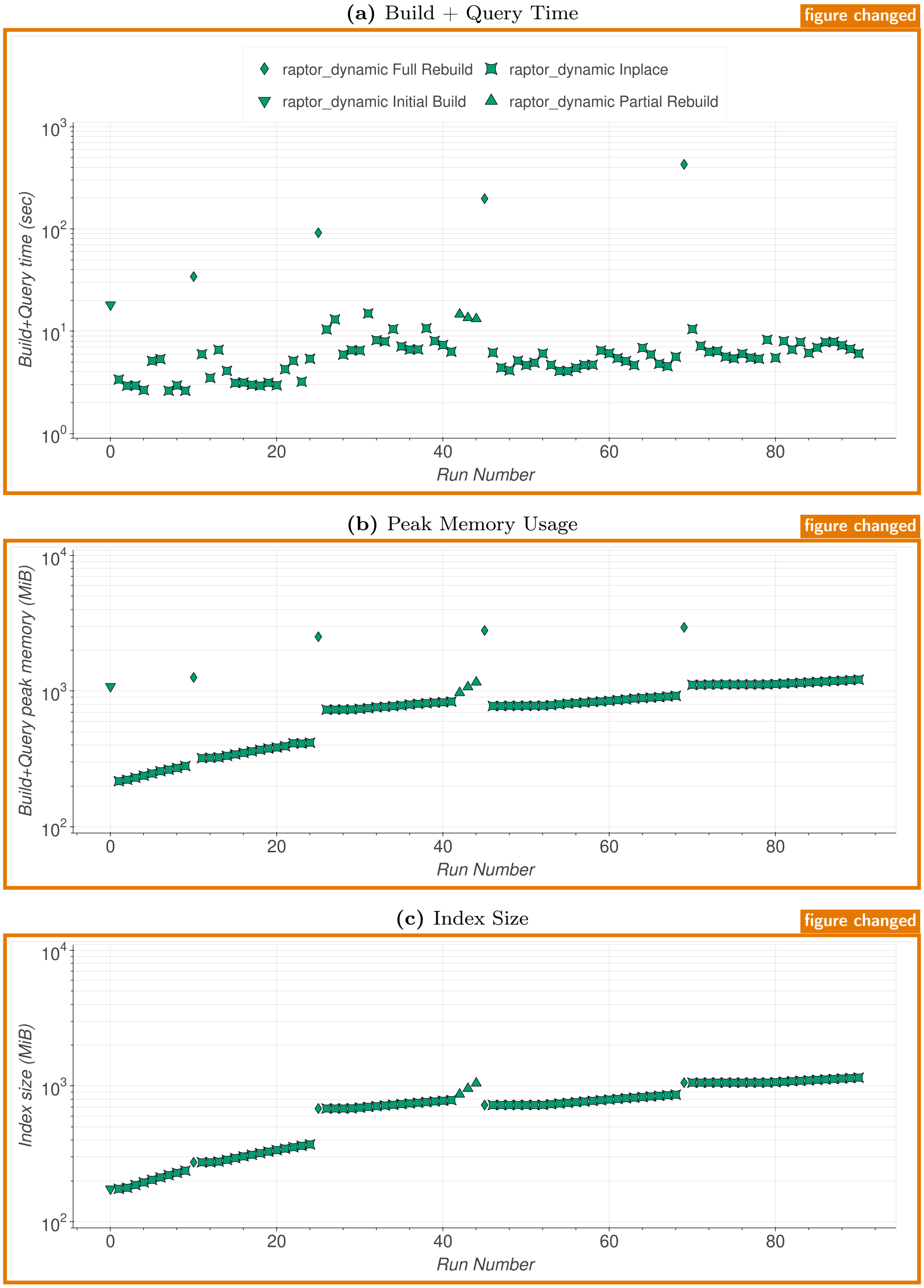


**Figure 9** **Batch Update with Insertions and Deletions of Evenly Simulated Dataset.** Build + Query time in seconds (a), peak memory usage in MiB (b), index size in MiB (c). The initial build (Run Number x=0) contains 10,000 sequences, and each incremental run (x>1) deletes 100 sequences and adds 1,000 sequences.

**(a)** Build Time

figure changed

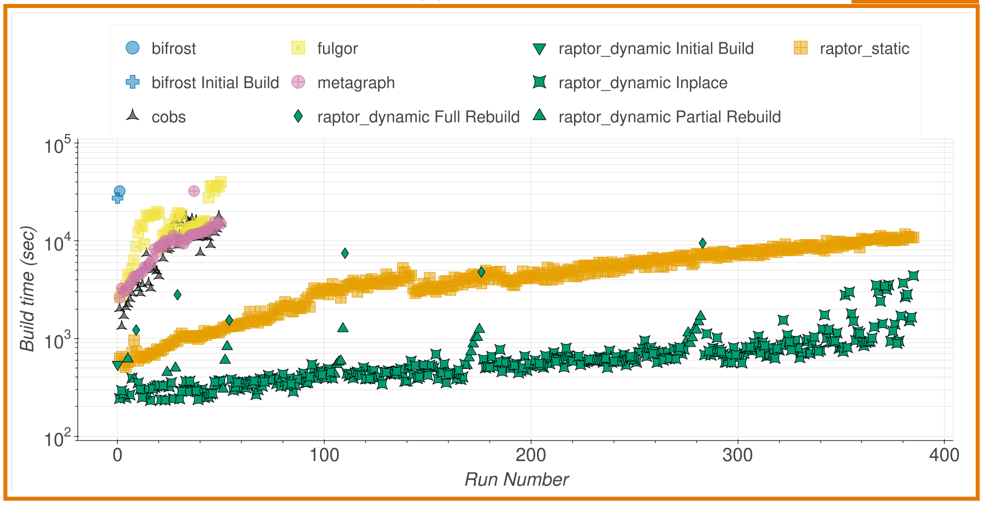


**(b)** Peak Memory Usage

figure changed

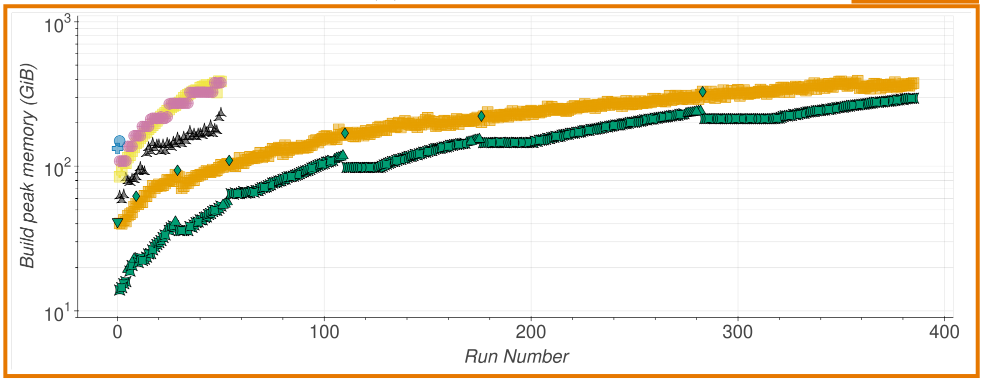


**(c)** Index Size

figure changed

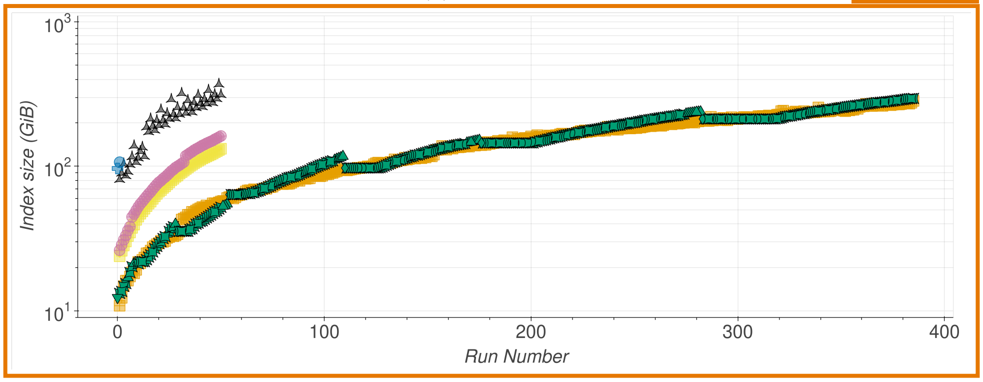


**Figure 10 Human RNA sub-sampled to 1%.** Build time in seconds (a), peak memory usage in GiB (b), and index size in GiB (c). The initial build (Run Number x=0~~, Raptor~~; Raptor-Dynamic and Bifrost only) contains 1,000 sequences, and each incremental run ($x \geq 1$) adds 100 sequences. COBS, Fulgor, and Metagraph were executed only for the first 50 batches due to long run times; Bifrost failed after the first update.

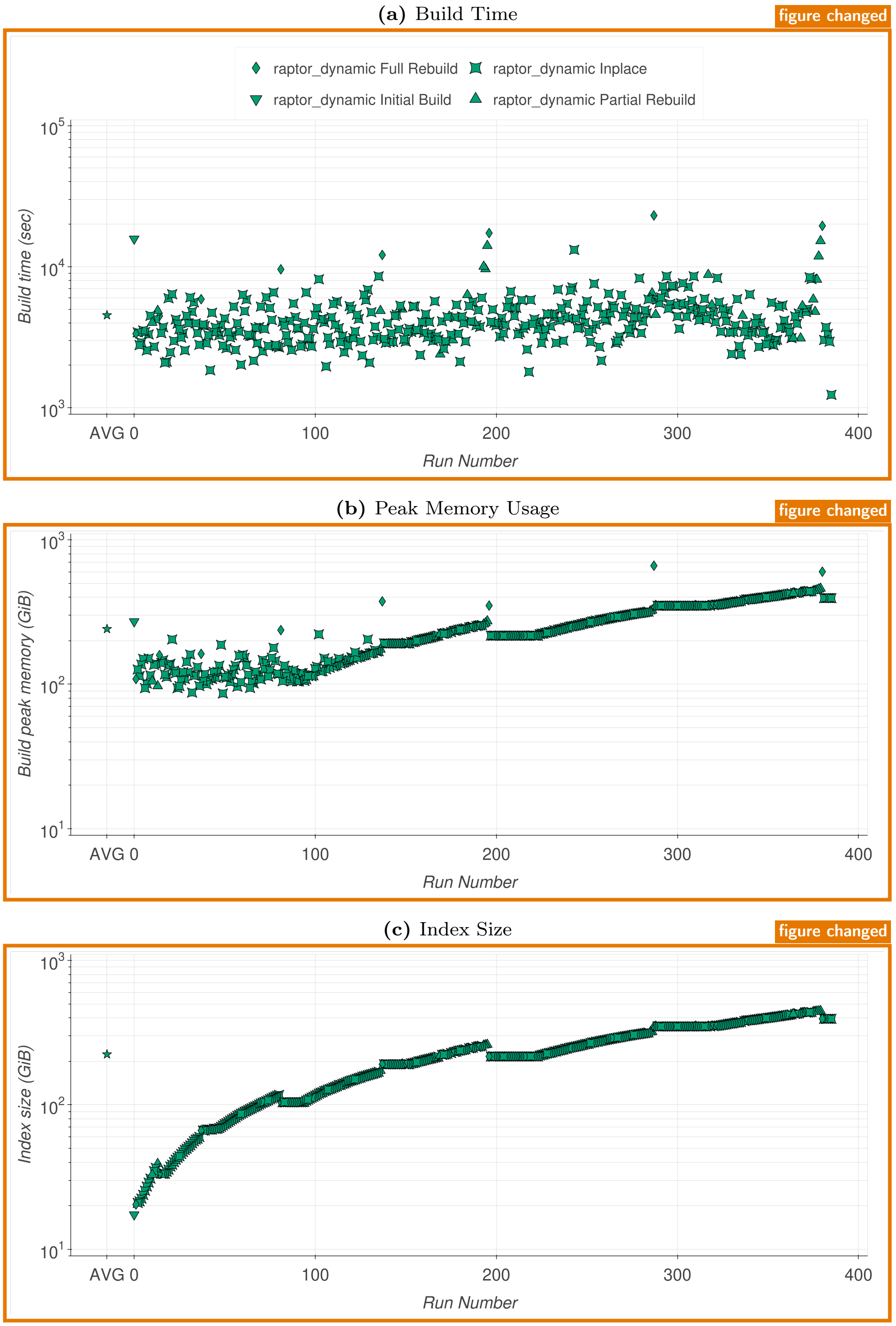


**Figure 11 Human RNA 39,400 full samples.** Build time in seconds (a), peak memory usage in GiB (b), and index size in GiB (c). The initial build (x=0) contains 1,000 sequences, and each incremental run (x≥1) adds 100 sequences. Low-frequency k-mers were removed based on file size.

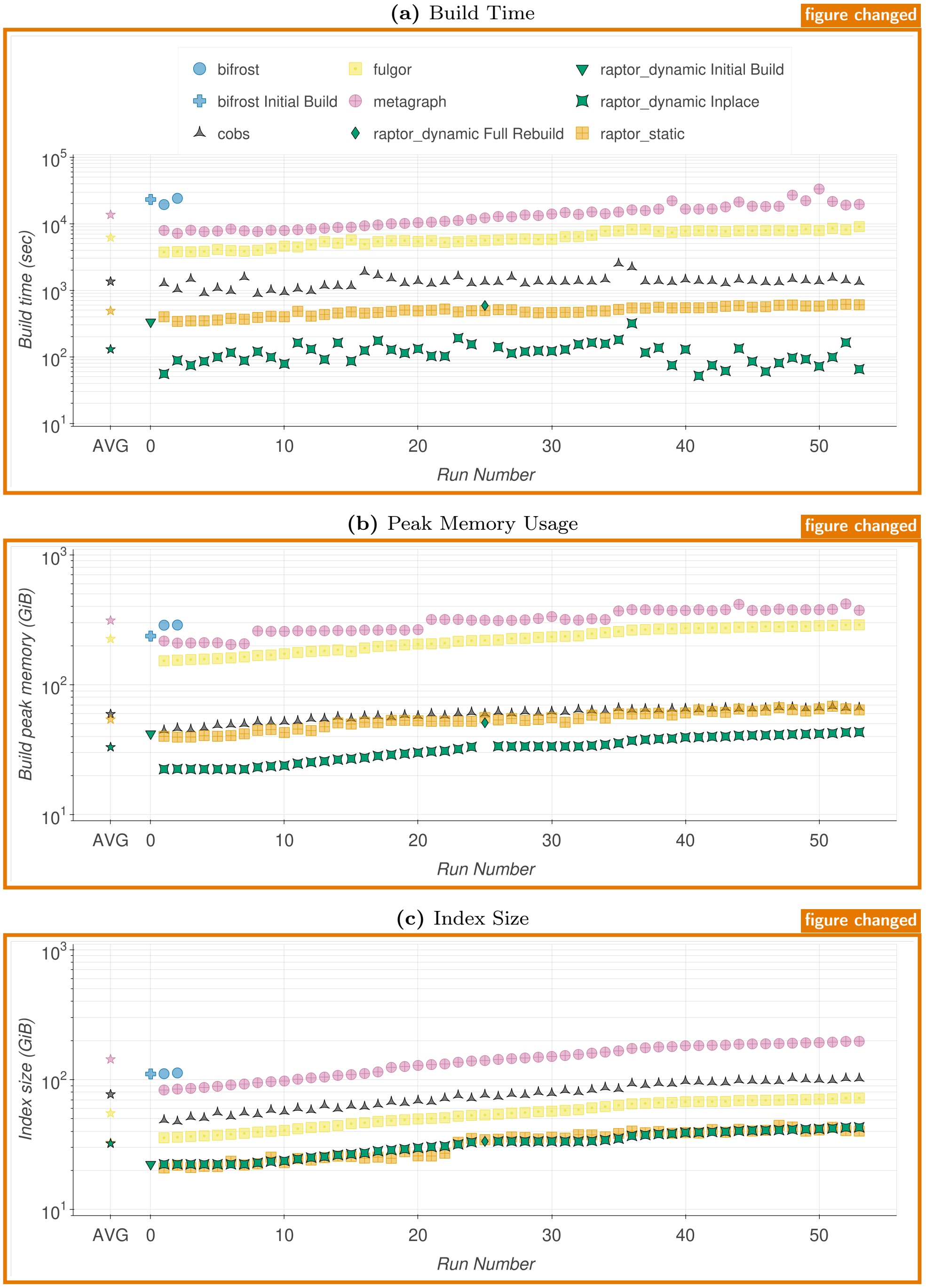


**Figure 12** **Weekly Batch Update of Real RefSeq Genomes.** Build time in seconds (a), peak memory usage in GiB (b), and index size in GiB (c). The initial build (Run Number x=0; Raptor-Dynamic and Bifrost only) contains 10,428 genomes, and each incremental run ($x \geq 1$) adds the genomes of one week. Bifrost failed after the second update.

#### Competing Interests
The authors declare that they have no competing interests.

#### Funding
Not applicable

#### Authors' contributions
KR initiated the project and the main idea for the DHIBF. MW devised in her MSc thesis a first prototype for a DHIBF. ES integrated the viable methods into the main HIBF branch. VP conducted the experiments and wrote the Results section. All authors read and approved the final manuscript.

#### Acknowledgements
Not applicable

#### Availability and Requirements
Project name: Raptor
Project home page: https://github.com/seqan/raptor
Operating system(s): Linux, macOS, BSD
Programming language: C++
Other requirements: For compiling: GCC $\geq$ 14, or Clang $\geq$ 19. No additional runtime requirements.
License: BSD-3-Clause
Any restrictions to use by non-academics: No restriction

#### Author details
[1]Department of Mathematics and Computer Science, Freie Universität Berlin, Takustr. 9, 14195, Berlin, Germany. [2]Max Planck Institute for Molecular Genetics, Ihnestraße 63-73, 14195, Berlin, Germany. [3]Julius Center for Health Sciences and Primary Care, University Medical Center Utrecht, Universiteitsweg 100, 3584 CG, Utrecht, The Netherlands.

### Appendix A: Appendix

#### A.1 Hardware specifications

*CPU*

```
Architecture:                          x86_64
CPU op-mode(s):                        32-bit, 64-bit
Address sizes:                         46 bits physical, 57 bits virtual
Byte Order:                            Little Endian
CPU(s):                                112
On-line CPU(s) list:                   0-111
Vendor ID:                             GenuineIntel
Model name:                            Intel(R) Xeon(R) Gold 6348 CPU @ 2.60GHz
CPU family:                            6
Model:                                 106
Thread(s) per core:                    2
Core(s) per socket:                    28
Socket(s):                             2
Stepping:                              6
BogoMIPS:                              5200,00
Flags:                                 avx avx2 [...]
                                       avx512{bw,cd,dq,f,ifma,vbmi,vl,_bitalg,_vbmi2,_vnni,_vpopcntdq}
Virtualization:                        VT-x
L1d cache:                             2,6 MiB (56 instances)
L1i cache:                             1,8 MiB (56 instances)
L2 cache:                              70 MiB (56 instances)
L3 cache:                              84 MiB (2 instances)
NUMA node(s):                          2
NUMA node0 CPU(s):                     [0,2,...,110]
NUMA node1 CPU(s):                     [1,3,...,111]
Vulnerability Gather data sampling:    Mitigation; Microcode
Vulnerability Indirect target selection: Mitigation; Aligned branch/return thunks
Vulnerability Itlb multihit:           Not affected
Vulnerability L1tf:                    Not affected
Vulnerability Mds:                     Not affected
Vulnerability Meltdown:                Not affected
Vulnerability Mmio stale data:         Mitigation; Clear CPU buffers; SMT vulnerable
Vulnerability Reg file data sampling:  Not affected
Vulnerability Retbleed:                Not affected
Vulnerability Spec rstack overflow:    Not affected
Vulnerability Spec store bypass:       Mitigation; Speculative Store Bypass disabled via prctl
Vulnerability Spectre v1:              Mitigation; usercopy/swapgs barriers and __user pointer sanitization
Vulnerability Spectre v2:              Mitigation; Enhanced / Automatic IBRS; IBPB conditional;
                                                   PBRSB-eIBRS SW sequence; BHI SW loop, KVM SW loop
Vulnerability Srbds:                   Not affected
Vulnerability Tsa:                     Not affected
Vulnerability Tsx async abort:         Not affected
Vulnerability Vmscape:                 Not affected
```

*Kernel*

```
Linux 6.1.0-43-amd64 #1 SMP PREEMPT_DYNAMIC Debian 6.1.162-1 (2026-02-08) x86_64 GNU/Linux
```

#### A.2 Supplemental Data

The source code is written in C++23 using the SeqAn Library [36] and available at https://github.com/seqan/raptor. Additional resources for this project, including sample metadata and analysis scripts, are available in the latter GitHub repository.

#### A.3 Existing tools

**Table 3 Algorithmic comparison of multi-set sequence indexing tools.** Overview of representative colored de Bruijn graph and sequence aggregation indices, detailing their support for dynamic insertion and deletion operations along with their core indexing mechanisms. Expanded version of Table 1.

| Tool | Ins | Del | Mechanism/Notes | Citation |
|---|---|---|---|---|
| Mantis (Classic/RRR) | × | × | Encodes k-mers using a Counting Quotient Filter (CQF) mapped to color IDs, which are stored as an RRR-compressed (static) bit matrix. | [22] |
| Dynamic Mantis (MST) | ✓ | × | Replaces RRR compression with an MST-based compression of color classes. Uses the Bentley-Saxe transformation to merge indexes iteratively. | [29] |
| Rainbowfish | × | × | Compresses graph colors by storing unique occurrence patterns in an optimized, minimal, static table. | [37] |
| Vari | × | × | Uses a BWT-encoded succinct de Bruijn graph (BOSS) combined with RRR and Elias-Fano compressed occurrence matrices. | [38] |
| Sequence Bloom Tree (SBT) | ✓ | ✓ | Organizes independent sample Bloom filters into a hierarchical binary tree where internal nodes store the union of their descendants. | [35] |
| Split Sequence Bloom Tree (SSBT) | ✓ | ✓ | Improves upon the SBT by splitting nodes into a "similarity filter" and a "remainder filter" to prune redundant lookups. | [39] |
| AllSome Sequence Bloom Tree (SBT-AS) | ✓ | ✓ | An SBT derivative categorizing bits as 'all' or 'some' at internal nodes to accelerate I/O and queries. Updates require recompressing RRR bitvectors. | [40] |
| BIGSI | ✓ | × | Stacks identical Bloom filters for each dataset into a column-major bitmatrix queried via a hash table. Index update requires modifying every key in the index. | [28] |
| COBS | × | × | A compact bit-sliced signature index linking an inverted index with Bloom filters, grouping files of similar sizes to save space. | [17] |
| Cortex | × | × | Original cdbg implementation storing k-mers and their edge linkages in a standard hash table. | [41] |
| VariMerge | ✓ | × | Extends the Vari architecture to support the efficient merging of distinct, updatable cdBG representations. | [27] |
| Bifrost | ✓ | ~ | Builds compacted colored de Bruijn graphs using a minimizer-based hash table mapped to unitigs. Deletion only via API. | [16] |
| SeqOthello | ✓ | × | Relies on Othello minimal perfect hashing to map k-mers to their corresponding occurrence profiles. Updates require a merge and hash reconstruction. | [26] |
| Bloom Filter Trie (BFT) | ✓ | ✓ | Encodes k-mer paths within a burst trie, using Bloom filters at compressed inner nodes for rapid lookups. | [42] |
| RAMBO | ✓ | × | Applies a Count-Min Sketch paradigm, randomly partitioning documents into repeated and merged Bloom filters. | [43] |
| HowDe-SBT | ✓ | ✓ | Optimizes SBTs by stripping out redundant "non-informative" bits, drastically shrinking the tree's memory footprint. | [19] |
| Raptor (IBF) | × | × | Stores representative k-mers in Interleaved Bloom Filters (IBFs). | [32] |
| HIBF | × | × | A Hierarchical IBF optimizing space by splitting large bins and merging small ones into multi-level interleaved filters. | [30] |
| PAC | ✓ | × | Partitions k-mers by minimizers, structuring Aggregated Bloom filters into Bloom Comb Trees. | [25] |
| MetaGraph (Static/Default) | × | × | Uses a BOSS-encoded SuccinctDBG and highly compressed annotation matrices (e.g., Multi-BRWT). | [24] |
| MetaGraph (SuccinctDBG Dynamic) | ✓ | ✓ | Adapts the static MetaGraph approach by backing the core graph vectors with dynamic, cache-efficient B-trees. | [24] |
| kmindex | ✓ | × | Builds partitioned, inverted one-hash Bloom filter matrices aligned by minimizers. | [20] |
| MetaProFi | ✓ | × | Uses a chunked Bloom filter matrix approach. Natively permits seamless index extension with new batches, but provides no mechanism for deletion. | [44] |
| Fulgor | × | × | Maps ccdBG unitigs to colors using an order-preserving SSHash dictionary, maintaining unitigs in exact color order to compress integer arrays. | [18] |

### A.4 Sequence Simulation

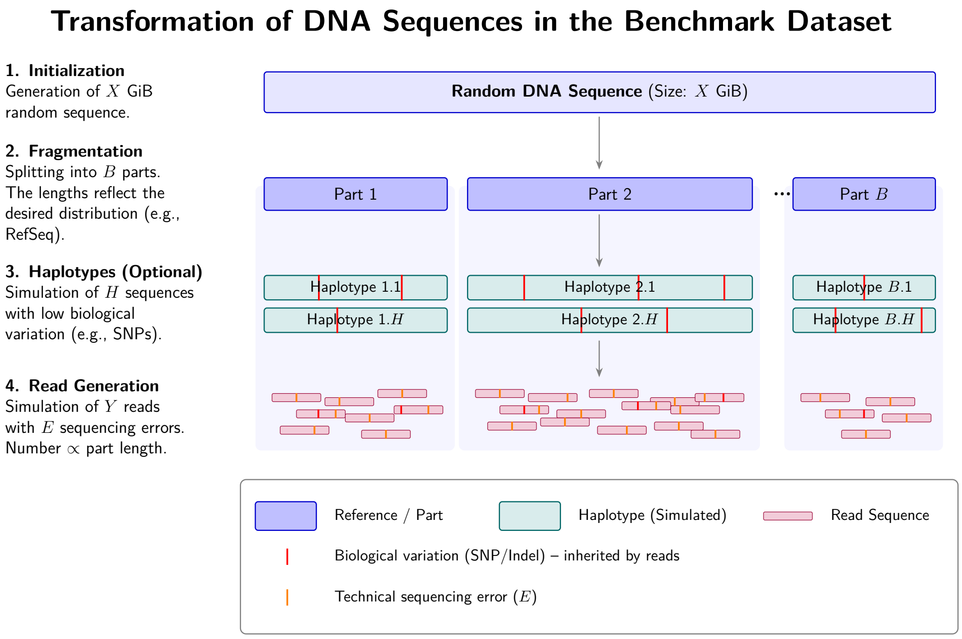


**Figure 13 Sequence Simulation.** From an initially generated random DNA sequence ($X$ GiB), $B$ parts are extracted via fragmentation. Optionally, $H$ haplotypes are simulated from these reference parts, containing controlled biological variations (e.g., SNPs/Indels, marked in red). Finally, read sequences are simulated from the haplotypes, with the number of reads scaling proportionally to the length of the respective part. The reads inherit the biological mutations and are additionally injected with simulated technical sequencing errors ($E$, marked in orange).

### A.5 Extensions

Splitting merged bins into multiple subtrees

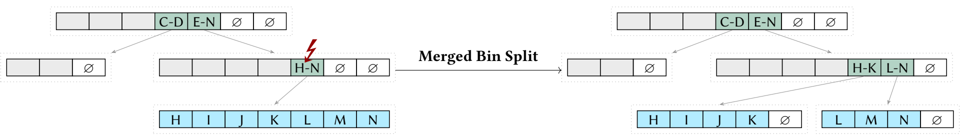


**Figure 14 Example merged bin split.** Empty bins are denoted by ∅. A gray background is used for present bins that are not relevant for the respective example. A merged bin exceeding the false positive threshold after sample insertion is marked with a ↯. Instead of a partial rebuild, the affected merged bin is split into two merged bins.
